# CADENCE: Open-source software for signal conditioning, feature extraction, visualization and analysis of cardiac optical mapping data

**DOI:** 10.64898/2026.09.30.755844

**Authors:** Kedar Aras, Shaivi Pandey, Adedayo Adelakun, Ethan Minnella, Alexis Lichtenberger, Neelam Qadus, Luke Barnhart

## Abstract

Optical mapping is a fundamental tool for cardiac electrophysiology research. Processing of these signals can be non-trivial, requiring multiple pre- and post-processing steps to limit interpretation errors. Existing software packages have features siloed across platforms and require users to combine several packages for in-depth analysis. To address this, we developed CADENCE, an open-source MATLAB-based application combining feature breadth (e.g., SVD denoising, conduction velocity, alternans, annotated rotors) and depth (e.g., 15 independent alternans features across both voltage and calcium modalities) in an end-to-end pipeline including data conversion, signal conditioning, feature extraction, and signal analysis. We demonstrate CADENCE’s capabilities across datasets spanning four species (rat, rabbit, pig, and human), three tissue preparations (slices, wedge, and whole heart), and three camera configurations (single, dual, and panoramic). CADENCE was validated against three blinded expert observers (15 hearts) and demonstrated agreement comparable to inter-observer variability (SD ratios 1.07/0.95/0.89 for APD80, activation, CaTD80). Dominant frequency was recovered with a median error of 0.03% across 84 of 88 paced recordings with 1:1 capture. CADENCE’s code is publicly available and distributed as a compiled standalone application that the end-user can install and run without a MATLAB license to encourage adoption and standardization across laboratories.

## INTRODUCTION

Optical mapping remains one of the fundamental tools for cardiac electrophysiology investigations into cardiovascular diseases^1^. Optical mapping experiments typically use voltage and/or calcium sensitive dyes to acquire high resolution spatiotemporal cardiac maps, which can provide an electrical profile of the heart under physiological and disease states^2^. These studies have advanced our understanding of pathologies such as ischemia^3^ and arrhythmias^4^, and enabled development of new therapeutic interventions^5,6^, thus highlighting the importance of optical mapping as a powerful cardiac electrophysiology research technique.

Processing of cardiac optical imaging data is non-trivial and can entail several pre-processing and post-processing steps to mitigate errors in data interpretation^7^. For example, raw fluorescence optical mapping data can include additive noise (e.g., 60 Hz), motion artifacts (e.g., working heart preparation or if the mechanical uncoupler is not sufficient to attenuate motion), and multiparametric signals (e.g., transmembrane voltage data, calcium transient data etc.). Consequently, analysis of optical mapping data requires pre-processing steps (e.g., spatial and temporal filtering, ensemble averaging) and post-processing steps (e.g., data masking, histogram thresholding etc.) to improve data quality.

Several software tools have been released over the years to help with the analysis of cardiac optical mapping data. Software packages including RHYTHM^8^, KairoSight^9^, and ElectroMap^10^ support data processing and analysis pipelines and have proven valuable to cardiac electrophysiology research labs, especially as many of them lack the computational expertise to build their own software. There are also other software packages available including ORCA^11^, COSMAS^12^, and Spiky^13^ that offer specialized but limited capabilities. For example, ORCA only supports CV measurements. COSMAS is script-only software and lacks support for arrhythmia analysis. Spiky does not support panoramic datasets. Consequently, many useful features are siloed across various platforms, requiring the use of multiple software packages to perform in-depth optical mapping data analysis. While labs often assemble scripts to help with data analysis, processing choices are often undocumented, and results are hard to compare between groups, amplifying reproducibility issues. For example, without a validated, documented pipeline, a measured APD or CV is not comparable across labs. A recent evaluation of KairoSight-3.0 and ElectroMap identified a lack of support for multi-parametric analysis, robust SNR qualification under arrhythmia, and comprehensive arrhythmia tooling^14^. Thus, there remains a need for a standardized open-source software platform for end-to-end optical mapping data analysis.

We introduce here CADENCE, an open-source MATLAB-based software toolkit with a modular, flexible architecture of five independently usable modules that supports features currently distributed across several packages (see **Table 1**), while also including capabilities (e.g., SVD denoising) not available in existing packages. CADENCE supports an end-to-end schema-validated pipeline—spanning data conversion (e.g., .TIFF to .mat), signal conditioning (e.g., spatial and temporal filtering, ensemble averaging), feature extraction (e.g., activation, APD, CaTD, alternans), and signal analysis (e.g., annotated arrhythmia rotor and wavefronts). While CADENCE combines feature breadth (e.g., conduction velocity) and depth (e.g., 15 independent alternans features across both voltage and calcium modalities), it currently lacks support for NADH/metabolic imaging and S1–S2 restitution analysis.

**Table 1.** CADENCE comparison: Feature comparison with existing software toolkits including ElectroMap, RHYTHM 4.0, and KairoSight 3.0. Y = Feature Available, N = Feature Unavailable, ∼ = Feature Availability Unclear. Other toolkits such as COSMAS, Spiky, and ORCA were not included in the comparison table due to their specialized but limited feature support.

| Feature | CADENCE<br>1.0 | ElectroMap | RHYTHM<br>4.0 | KairoSight<br>3.0 |
| --- | --- | --- | --- | --- |
| **Platform & Deployment** |  |  |  |  |
| Language | MATLAB | MATLAB | MATLAB | Python 3.8 |
| Runs without MATLAB license | Y | Y | N | Y |
| Open source | Y | Y | Y | Y |
| Batch / high-throughput processing | Y | Y | Y | Y |
| Schema validation | Y | N | N | N |
| **Data Input Support** |  |  |  |  |
| SciMedia MiCAM (.gsd) | Y | Y | Y | N |
| TIFF input (.tiff) | Y | Y | N | Y |
| Dual voltage–calcium | Y | Y | Y | Y |
| NADH / FAD imaging | N | N | Y | N |
| Panoramic / multi-camera | Y | N | Y | N |
| **Signal Processing** |  |  |  |  |
| SVD denoising | Y | N | N | N |
| Spatial binning | Y | Y | Y | Y |
| Temporal filtering | Y | Y | Y | Y |
| Drift correction | Y | Y | Y | Y |
| Data normalization | Y | Y | Y | Y |
| Ensemble averaging | Y | Y | Y | Y |
| Image alignment & registration | Y | N | N | Y |
| Data mask (user mask, SNR, histogram) | Y | Y | Y | Y |
| Adaptive SNR masking | Y | N | N | N |
| Motion correction | Y | N | N | N |
| **Electrophysiology Feature Extraction** |  |  |  |  |
| Activation / Rise time | Y | Y | Y | Y |
| Conduction velocity | Y | Y | Y | Y |
| APD / Repolarization | Y | Y | Y | Y |
| Calcium transient duration | Y | Y | Y | Y |
| Calcium decay constant | Y | Y | Y | Y |
| Voltage–calcium coupling | Y | Y | Y | Y |
| **Arrhythmia / complexity analysis** |  |  |  |  |
| DF/RI/OI | Y | ~ | ~ | ~ |
| Alternans | Y | Y | Y | Y |
| Phase maps | Y | Y | Y | N |
| Annotated rotor and wavefront tracking | Y | N | N | N |

In this study, we demonstrate CADENCE’s capabilities across optical mapping datasets spanning four species (rat, rabbit, pig, and human) with a range of cycle lengths (85–1000 ms), three tissue preparations (slices, wedge, and whole heart), and three camera configurations (single, dual, and panoramic). We also validate CADENCE against blinded manual marking by three observers and against the pacing stimulus for dominant frequency (DF). CADENCE’s code is publicly available (MIT license) and distributed as a compiled standalone application that the end-user can install and run without a MATLAB license.

## METHODS

We provide a high level summary of the experiments and study protocols used for data acquisition, and computational approach for signal conditioning and feature extraction algorithms. Detailed description (e.g., animal characteristics, anesthesia, dye loading) can be found in the Supplementary Methods.

### Data Acquisition

All studies using animal models including pig, rabbits, and rats are in accordance with the ARRIVE guidelines 2.0 and approved by the Institutional Animal Care and Use Committee respectively at University at Buffalo (Protocol Approval Numbers: 202200063, 202200066, 202300053). Use of de-identified human heart tissue (Connect Life, Buffalo, NY) was approved by Institutional Review Board at University at Buffalo (Protocol Approval Number: 6939). All procedures followed the National Institutes of Health Guide for the Care and Use of Laboratory Animals.

To validate CADENCE software across a variety of optical imaging datasets, the experimental setups included single camera (V_m_ or Ca), dual camera (V_m_ and Ca) and panoramic (four cameras) configuration. We also employed three different tissue preparations including: cardiac slices (pig), left ventricular (LV) wedge (donor human hearts), and whole heart (rat and rabbit). Tissue was immobilized with blebbistatin to suppress motion artifacts in optical recordings, without adverse electrophysiological effects^4,8,15–17^. Signals were recorded using up to four MiCAM03/05 (SciMedia, CA) CMOS cameras (256 x 256 pixels or 100 x 100 pixels, 1 kHz sampling rate). Study protocol entailed conducting standard S1-S1 pacing restitution protocol at decreasing pacing cycle length until loss of 1:1 capture. We then administered isoproterenol to shorten action potential duration. Arrhythmia was then pace-induced using S1-S1 protocol or 1-second burst pacing at 60 Hz.

### Software Architecture and Algorithms

**Fig. 1** depicts the CADENCE software architecture that follows a modular design philosophy with independently usable modules including for data conversion, signal conditioning, feature extraction, signal analysis, and conduction velocity. Graphical user interfaces (GUI) for each module were created using MATLAB App Designer, which employs the object-oriented coding framework (see **Supplementary Fig. S1)**.

**Fig. 1.**
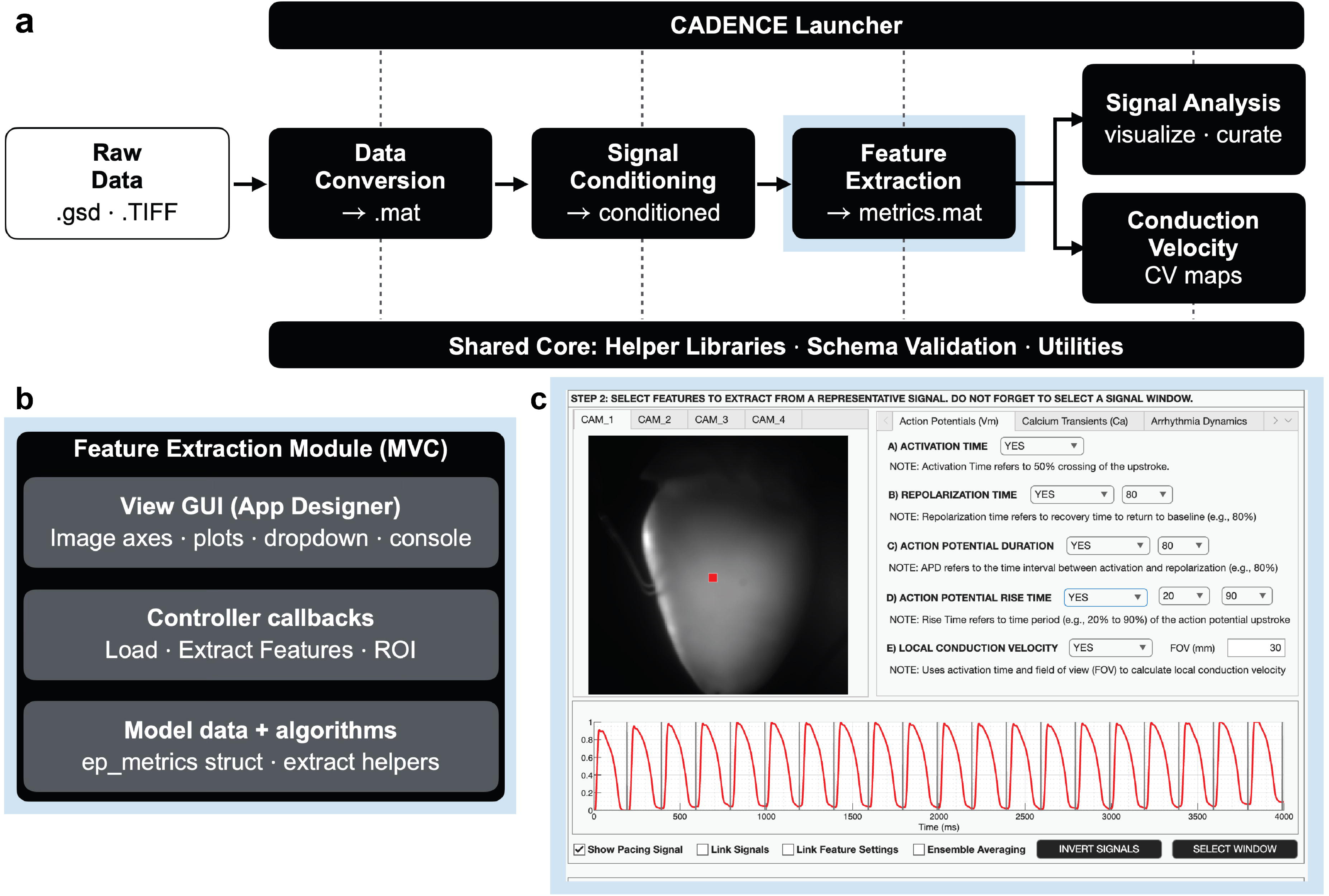
CADENCE Software Architecture: **a)** CADENCE software employs a modular approach to building the data processing and analysis pipeline (data conversion, signal conditioning, feature extraction, signal analysis, and conduction velocity) with shared core libraries. **b)** Feature Extraction module demonstrating the MATLAB supported Model-View-Controller (MVC) design pattern that CADENCE employs to build a flexible and extensible software toolkit. **c)** CADENCE user interface for the Feature Extraction module.

CADENCE supports TIFF and SciMedia format of raw data files for conversion to MATLAB file format. The signal conditioning pipeline comprises, in order: generating SNR map, drift correction, spatiotemporal denoising using singular value decomposition based low-rank decomposition ^18^ (falling back to 5×5 binning on any camera SVD cannot be applied to), spatial binning, temporal filtering, motion artifact correction^19,20^, normalization, polarity check, ensemble averaging^7^, and SNR mask (fixed threshold SNR ≥2). While motion correction algorithm is implemented, we did not demonstrate it here as our data acquisition experiments used a mechanical uncoupler.

Feature extraction includes support for voltage and calcium metrics including activation, repolarization, action potential duration (APD), action potential rise time, and local conduction velocity, V_m_-Ca delay, Ca decay time, Ca transient duration, Ca decay constant, and Ca rise time. For dual mapping, CADENCE supports image registration using rigid alignment of V_m_-Ca signals. Activation time is computed as sub-frame 50% upstroke. Sub-frame interpolation removes the quantization of frame-limited timing and improves the accuracy of the downstream conduction velocity calculation. The above metrics are computed using the ensemble-averaged beat unless the user selects a representative beat for feature extraction. Adaptive SNR masking is a key feature of feature extraction and uses the rule SNR threshold = max (1.5, 0.5 x median SNR of pixels with SNR > 1.5). CADENCE uses the full beat series to extract 15 alternans (voltage and calcium) metrics, and arrhythmia metrics (dominant frequency, regularity index, organization index, wavefront dynamics, and rotor dynamics). DF uses the rate adaptive search band and parabolic sub-bin peak refinement.

The conduction velocity (CV) module supports the computation of longitudinal (L) and transverse (T) CV from the local activation time map. By default, CV is computed using the inverse-gradient estimator with a local polynomial gradient fit. Field of view varies by species (e.g., rat 20 mm, human wedge 50 mm) and scales CV linearly. The longitudinal and transverse CV are computed using velocity-vector ellipse fit. Importantly, L/T is not reported if the activation map is planar (R^2^ > 0.8).

### CADENCE Validation

We employed blinded manual marking by three observers (15 hearts including 4 human, 4 rat, 4 pig, and 3 rabbit) for CADENCE vs human and human vs human validation. Briefly, the validation protocol comprised the overlap (4 recordings marked by all 3 observers) and breadth (each observer marking additional 4 recordings) arms. For each recording, the observer marked diastolic baseline, activation time (50% crossing), peak, repolarization (80% crossing) for 100 curated pixels (signals deemed to be of good enough quality by the observer to successfully mark the four locations). The extracted metrics (activation, APD80, and CaTD80) were compared against CADENCE. In addition, 88 recordings (from 83 hearts) were used to recover DF and compared against pacing stimulus (84 recordings had 1:1 capture).

### Use of AI-Assisted Technologies

Claude Opus 5.5 (Anthropic) was used to assist with code review, debugging, documentation, verification and fact checking. No AI tool was used to generate any images or figures. All AI-suggested code changes were tested and verified by the authors, who take full responsibility for the content of this manuscript.

## RESULTS

CADENCE’s code is publicly available (MIT license) and distributed as a compiled standalone application (https://github.com/kedararas/Cadence) that the end-user can install and run without a MATLAB license. Here, we demonstrate some of CADENCE capabilities for processing and analysis of optical mapping data.

### Signal Conditioning

**Fig. 2** displays the stages of processing raw signals including voltage (**Fig. 2a**) and calcium (**Fig. 2b**) from a representative rat optical mapping recording (paced at 175 ms). The sample raw traces are subjected to drift correction, spatiotemporal filtering (SVD denoising), temporal filtering (Butterworth [0 50 Hz]), normalization, and ensemble averaging (of 22 beats) to get cleaner signals with amplified SNR. Consequently, the SNR improves for the voltage trace from 5.3 to 76.3 and calcium trace from 10.0 to 82.3, with SVD spatiotemporal denoising contributing the largest increase. SNR in this instance refers to per-beat (peak-to-peak of the beat average divided by RMS deviation about it) and is distinct from the pipeline’s SNR mask, which depends on the recording length.

**Fig. 2.**
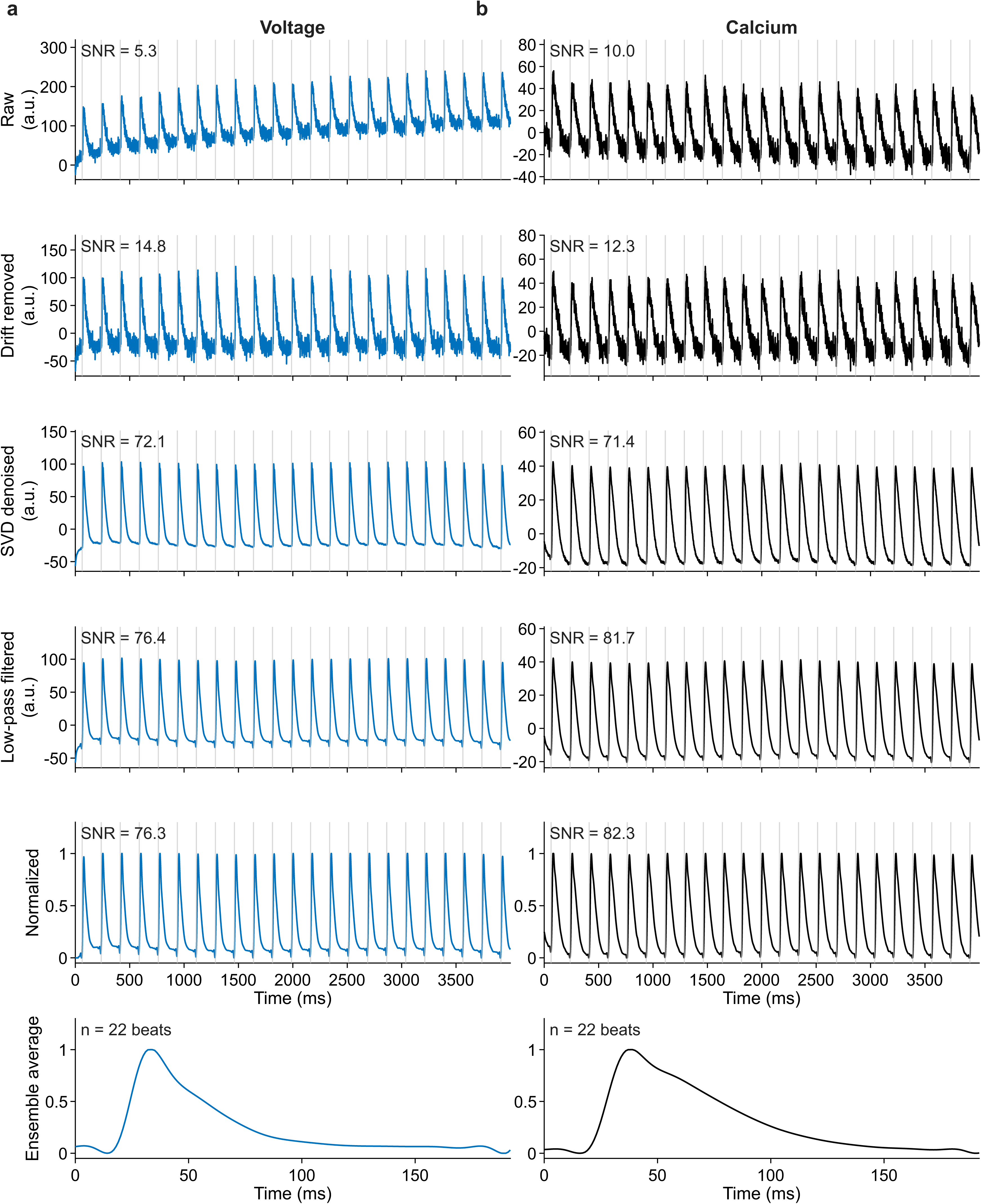
Signal Conditioning: Representative **a)** voltage and **b)** calcium traces (from one pixel location) from a rat whole heart setup paced at 175 ms subjected to the signal conditioning pipeline including drift removal, SVD spatiotemporal denoising, temporal filtering (50 Hz Butterworth low pass filter), normalization, and ensemble averaging steps, along with the individual effects on SNR.

### Feature Extraction

**Fig. 3** demonstrates sample metrics (activation time and action potential duration) extracted from different optical mapping datasets visualized as 2D cardiac maps as well as overlaid on representative traces. **Fig. 3a** shows activation and APD80 data extracted from a pig cardiac slice, single camera (voltage) optical mapping data recording paced at 300 ms. Median APD80 is 201.7 ms (IQR 199.0 – 204.0 ms, number of pixels = 52,392). The start and the end of the red overlay on the representative trace marks the instance of activation and repolarization (80%) at that pixel location. **Fig. 3b** highlights similar maps extracted from a rat whole heart Langendorff dual (voltage and calcium) camera optical mapping data recording paced at 175 ms. In addition to activation and APD80 maps, we also show activation delay (V_m_-Ca delay), and calcium transient duration (CaTD80) maps corresponding to the calcium traces. Median APD80 (43.6 ms with IQR: 39.6-49.4 ms), CaTD80 (77.9 ms), and V_m_-Ca delay (4.97 ms) are also shown. **Fig. 3c** demonstrates CADENCE support for four camera (panoramic) data sets. Activation and APD80 maps were generated from a rabbit whole heart Langendorff (voltage) optical mapping data recording paced at 200 ms. The rabbit median APD80 across the 4 cameras was in the range 123.4-125.9 ms. Importantly, the cross camera agreement was within 2.5 ms.

**Fig. 3.**
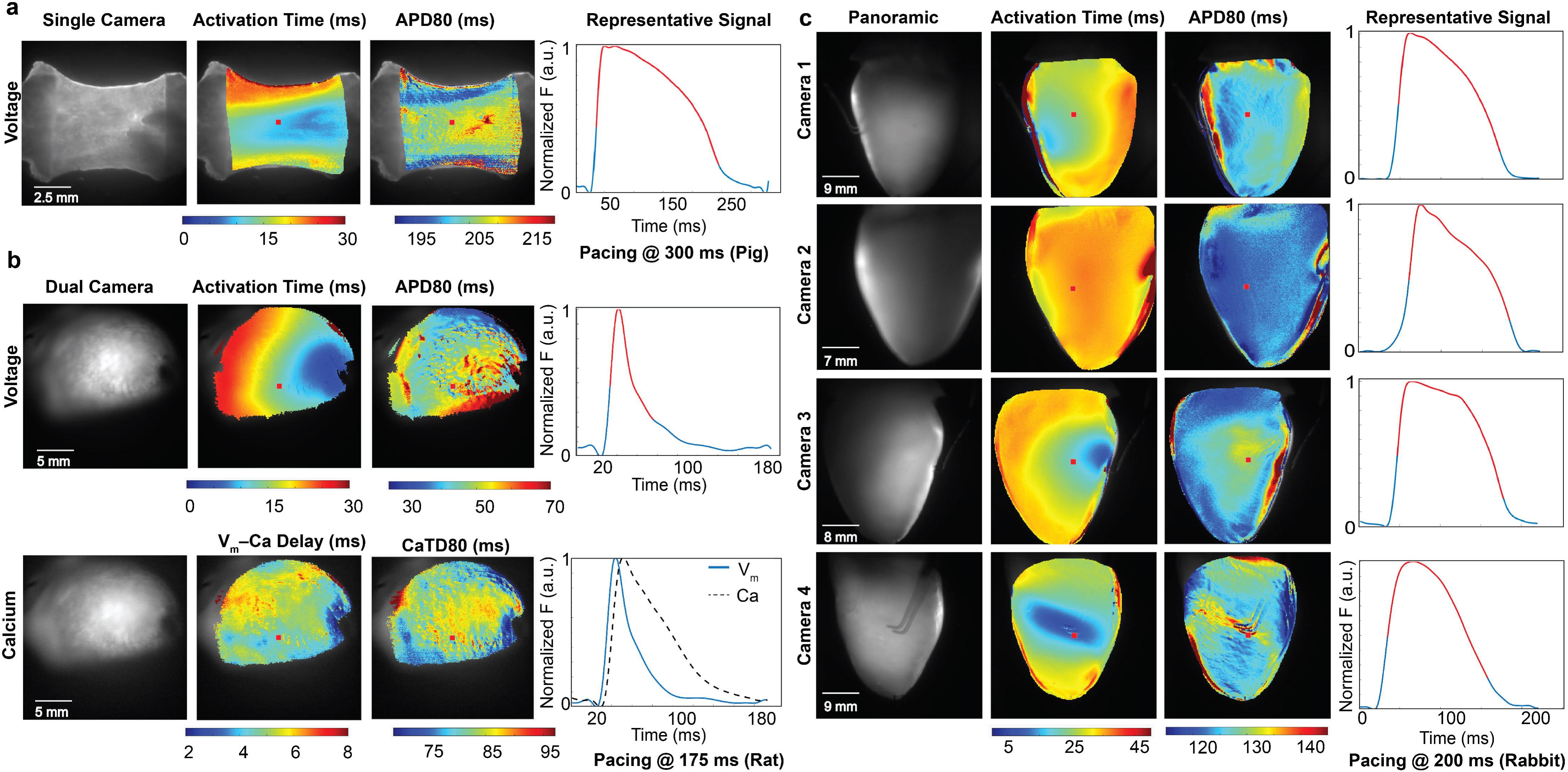
Feature Extraction: Representative metrics extracted from different species (pig, rat, rabbit) and tissue preparations (cardiac slice, Langendorff whole heart). **a)** Activation time, APD80 and trace from pixel location (red) from a single camera pig cardiac slice paced at 300 ms. The trace overlaid (red) represents the interval between the activation time and repolarization time (80%) and reflects the computed APD80 at that location. **b)** Activation time, APD80, V_m_-Ca activation delay, and CaTD80 from a dual mapping (two cameras) rat whole heart setup, paced at 175 ms and the corresponding traces. **c)** Activation time, APD80 and traces from panoramic (four cameras) rabbit whole heart setup, paced at 200 ms.

### Alternans Analysis

**Fig. 4** highlights CADENCE support for voltage (**Fig. 4a**) and calcium (**Fig. 4b**) alternans analysis using a rat dual camera (voltage and calcium) optical mapping data recording paced at 85 ms (47 beats and 23 pairs). The metrics displayed include cardiac maps for action potential (AP) amplitude ratio, AP alternans phase, APD50 alternans ratio, calcium transient (CaT) amplitude ratio, CaT alternans phase, and Ca diastolic alternans ratio as well as the representative traces (**Fig. 4c**). AP amplitude ratio (0.116) represents the beat-pair difference in AP amplitude (V_max_ –V_rest_) and provides a full tissue reference map but can be sensitive to signal artifacts and should be interpreted within the context of SNR. AP phase represents alternans phase map with sign of APD50 alternans (+1 odd beat is longer APD50, -1 even beat is longer APD50) highlighting in phase and anti-phase (discordant) alternans regions^21^. APD50 alternans ratio (0.172) represents the beat-pair difference in APD normalized to mean APD per pixel. It represents the canonical repolarization alternans and primary dispersion substrate. Similarly, CaT amplitude ratio (0.486) represents the beat-pair difference in Ca-transient amplitude (peak -onset) and is the canonical calcium alternans metric, highlighting the SR calcium cycling instability. Ca alternans phase represents the sign and continuous angle of Ca amplitude alternans. Ca alternans phase can be multiplied by AP phase map to produce Ca-Vm concordance map, with values of +1 representing in phase, and -1 describing out of phase (discordant) regions. Ca diastolic alternans ratio (0.240) represents the diastolic load instability. When diastolic alternans ratio is approximately 0, the alternans can be construed as being release driven alternans^22^. There are additional alternans metrics available as part of the CADENCE feature extraction module and documented in the supplementary materials.

**Fig. 4.**
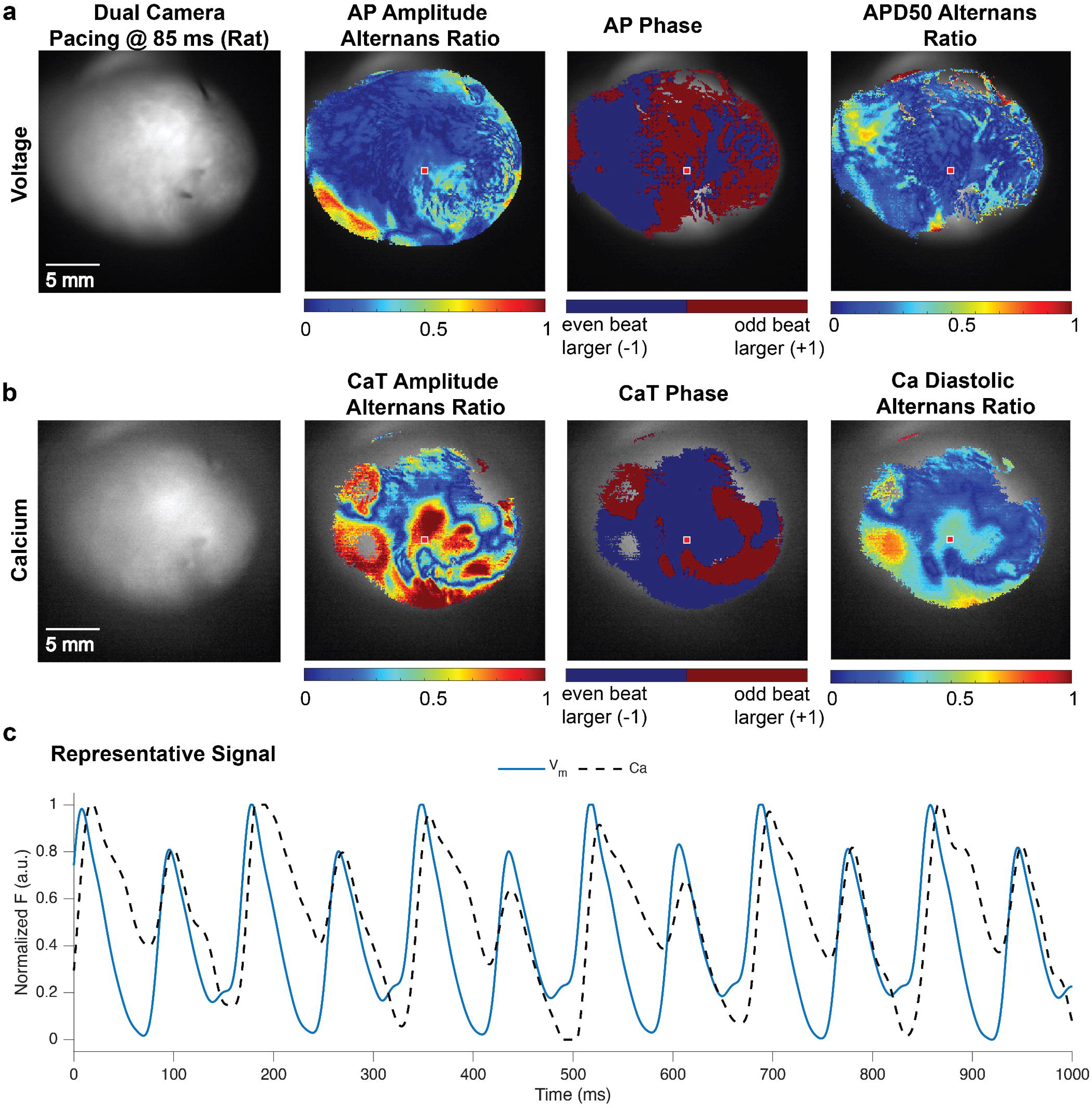
Alternans analysis: Voltage and calcium alternans from a rat whole heart preparation (paced at 85 ms). **a)** Voltage alternans metrics including AP amplitude alternans ratio, AP alternans phase (concordant/discordant) map, APD50 alternans ratio. **b)** Calcium alternans metrics including CaT amplitude alternans ratio, CaT alternans phase map, Ca diastolic alternans ratio. **c)** Traces (V_m_ and Ca) from the pixel location (red).

### Arrhythmia Analysis

**Fig. 5** shows an example of monomorphic ventricular arrhythmia using a rat optical mapping data recording induced using S1-S1 pacing protocol. **Fig. 5a** shows dominant frequency (DF) at 25.15 Hz (number of pixels = 37,115), regularity index (RI) at 0.651 (IQR 0.433-0.822), and Organization Index (OI) at 0.661 (IQR 0.440-0.838) characterizing the arrhythmia activity^23,24^. DF can be used for arrhythmia driver location, as the fastest activating region typically sources the rotor or focal driver^25,26^. High RI and OI values (RI →1, OI →1) represent regular and organized activity, as is the case with monomorphic arrhythmia seen in this example. Alternatively, Low RI and OI values (RI → 0, OI →0) suggest disorganized and chaotic activity, typically seen during atrial and ventricular fibrillation. CADENCE also supports visualization of arrhythmia activity in voltage and phase domain as well as automated annotation of wavefronts and stable rotors (defined as lasting at least 1.5 rotations) as seen in **Fig. 5b**. The trace window (3000 ms – 4000 ms) and the highlighted time window (in red) is visualized in both voltage and phase maps. The rotor can be tracked over the 40 ms interval completing one rotation (see **Supplementary Video S1**).

**Fig. 5.**
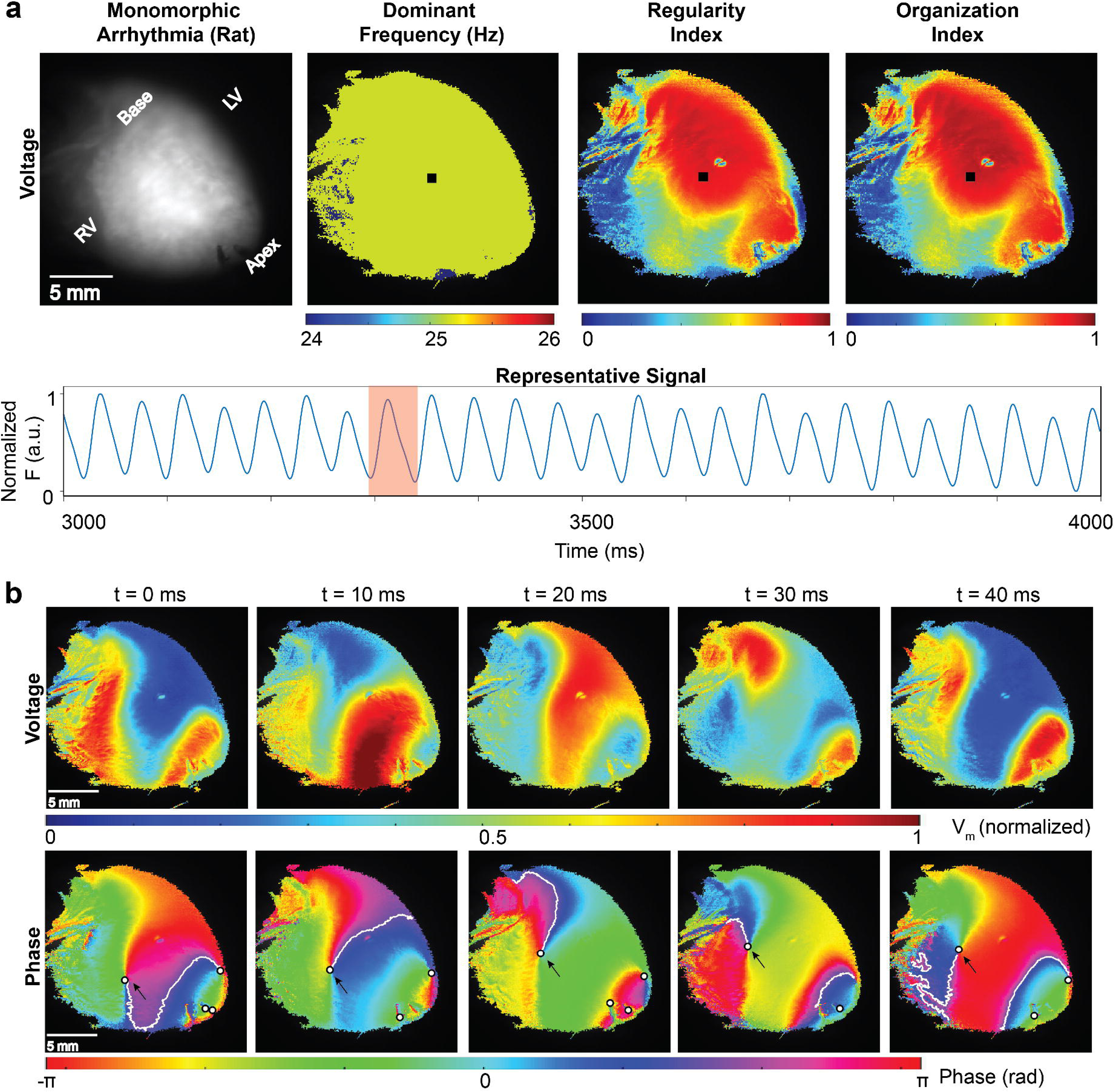
Arrhythmia analysis: Monomorphic ventricular arrhythmia from a rat whole heart preparation shows **a)** Background image with scale bar and anatomical orientation, Dominant Frequency (DF), Regularity Index (RI), and Organization Index (OI) cardiac maps with representative per-pixel normalized trace (black square on the maps) and overlaid 40 ms time window (red rectangle). The RI and OI look identical, which is expected for a near-sinusoidal rhythm with little harmonic power (also affected by the 50 Hz temporal filter cutoff). **b)** Voltage maps (normalized per pixel fractional depolarization) and annotated phase maps (Hilbert) corresponding to the time window and shown at 10 ms interval corresponding to one complete DF cycle. One of the stable rotors (white center marked by black outline) is tracked (black arrow) as it completes one rotation. The white contour lines represent the segmented wavefronts (and not a phase isoline).

### Conduction Velocity Analysis

**Fig. 6** demonstrates the CADENCE capability of computing conduction velocities from human LV wedge paced at 1000 ms (**Fig. 6a**) (FOV: 50 mm, 100 x100 grid, pixel pitch: 0.5 mm/pixel) and rat whole heart paced at 175 ms (**Fig. 6b**) (FOV: 20mm, 256 x 256 grid, pixel pitch: 0.078 mm/pixel) setups. Each panel shows the background image (with scale bar), activation and local CV maps, and histogram of CV distribution in the fitted region of interest (ROI). CV were computed using the same configuration including average vector (ellipse fit) method, polyfit engine, and auto-detection of pacing site. Both preparations show a focal breakthrough with concentric isochrones (non-planar) and pass the criteria for ellipse-fit (R^2^ > 0.3) at 0.81 and 0.88 respectively and planar-fit (R^2^ < 0.8) at 0.03 and 0.56 respectively as shown in table. The anisotropy ratio for both preparations exceeded the 1.5 threshold at 2.71 (L = 83.7 cm/s, T = 30.9 cm/s) and 2.21 (L = 76.3 cm/s and T = 34.5 cm/s) and were comparable even with 6.4x difference in pixel pitch (0.5 mm/pixel vs 0.078 mm/pixel) suggesting internal consistency in CV calculations. The fiber axes (30° vs 175°) represent the fast axis (longitudinal CV direction shown by fitted black L arrow) and reflect the tissue orientation in the field of view but has no other physiological significance. The difference in the number of valid pixels in the fitted ROI across the two preparations are reflective of the grid resolution (100 x 100 grid vs 256 x 256 grid). As can be seen from the CV histogram distribution the median CV values are 48.0 cm/s and 44.3 cm/s respectively over the fitted ROI with distribution peaking around 35-40 cm/s. The activation maps color ranges are different on account of different preparation sizes and pacing cycle lengths.

**Fig. 6.**
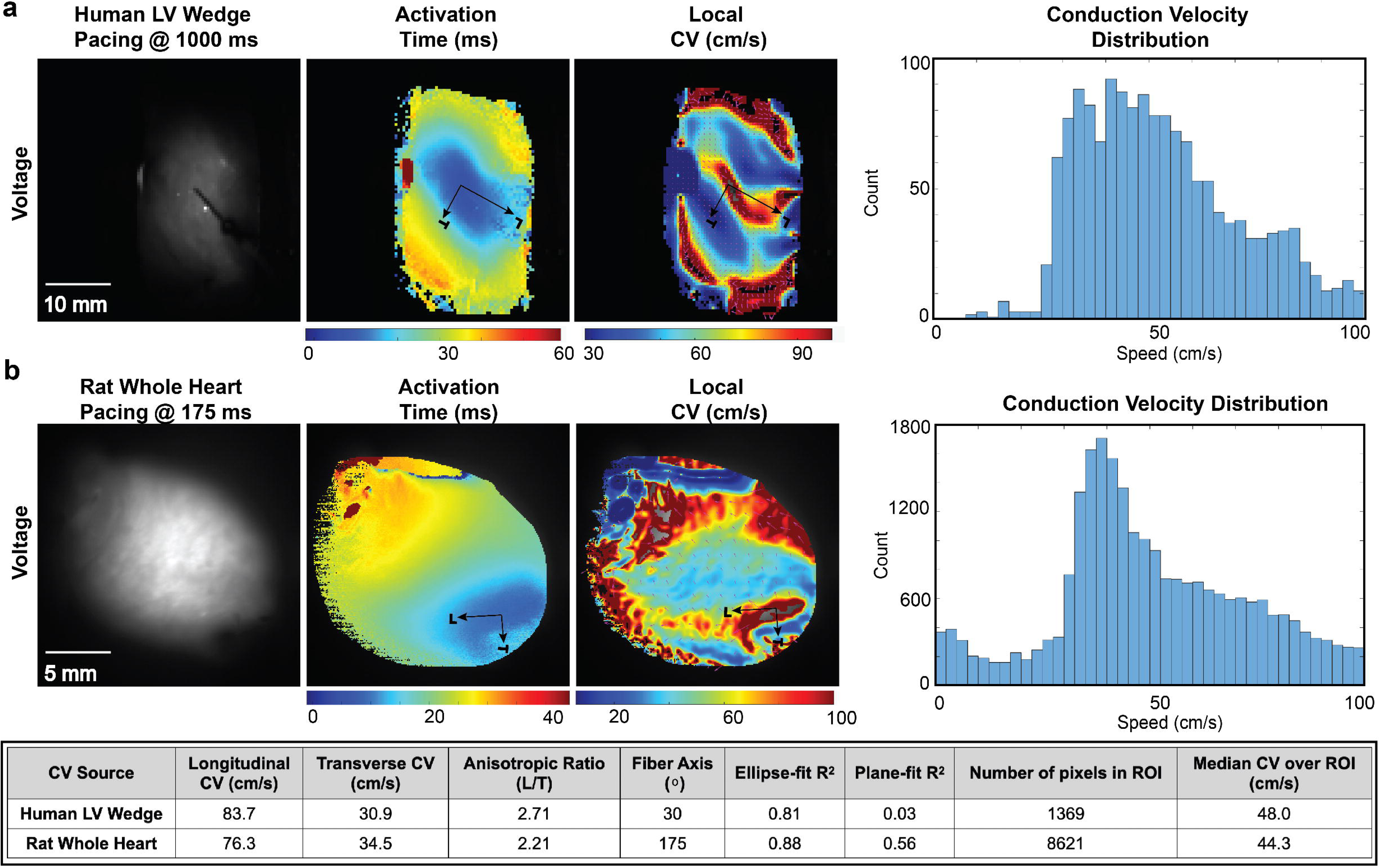
Conduction Velocity (CV) analysis: Background image with scale bar for field of view (FOV), activation map (ms), local CV map (cm/s), and histogram of CV distribution (cm/s) within the region of interest (ROI) from **a)** Human left ventricle wedge preparation (100 × 100 pixel grid, 50 mm FOV, pixel pitch 0.5 mm/pixel) at 1000 ms epicardial pacing and **b)** Rat whole heart preparation (256 × 256 pixel grid, 20 mm FOV, pixel pitch 0.078 mm/pixel) at 175 ms epicardial pacing. The computed CV values and validation parameters including longitudinal CV, transverse CV, anisotropic ratio, fiber axis (along longitudinal CV direction), ellipse-fit R^2^, plane-fit R^2^, number of pixels in ROI, and median CV within the fitted region (ROI) are reported in the table. CV was calculated using the following parameters: CV method: average vector, CV engine: Polyfit, Pacing site: Auto-detected. The L and T arrows mark the fitted fiber axis and its perpendicular. At point pacing, a low plane-fit R^2^ is expected.

### CADENCE Validation

**Fig. 7** demonstrates CADENCE vs human and human vs human validation using blinded manual marking by three observers (15 hearts/15 recordings including 4 human, 4 rat, 4 pig, and 3 rabbit). **Fig. 7a** shows mean APD80 map (interpolated) of the three observers compared with CADENCE APD80 map (non-interpolated) from an overlap human LV wedge recording (paced at 1000 ms). **Fig. 7b** shows the per pixel differences for the same APD80 pixel locations between CADENCE and the lead observer (who built the pixel pool) as well as between observer 2 and the lead observer. CADENCE vs human performance was SD 7.3 ms vs SD 5.3 ms with shared reference observer, so the spread compares directly. **Fig. 7c** shows the plot of Bland-Altman analysis with replicate measurements per subject (heart) without repeated measures correction. The pixel pairs (2382 pairs, 15 recordings, 3 observers) are not independent, and the pooled statistics are descriptive. The 95% Limits of agreement (LoA) at -9.2 /6.2 ms are centered on -1.5 ms. The grey band is the human vs human LoA. There were 7 pairs beyond the ±25 ms drawn at the edge. The SD ratios (CADENCE vs human divided by human vs human) were for APD80, Activation time, and CaTD80 at 1.07, 0.95, and 0.89 respectively (**Supplementary Table S1**). **Fig. 7d** shows the plot of agreement vs tolerance (at 5 ms) between CADENCE vs human and human vs human for the marked metrics including APD80 (92.9% vs 91.8%), Activation time (98.8% vs 99.1%), and CaTD80 (94.0% vs 97.2%). The lower plot uses the tolerance relative to the pair mean for APD80 and CaTD80 metrics. Both plots represent the overlap arm. **Fig. 7e** shows the median absolute CV error (%) vs activation time jitter (noise) computed from synthetic activation maps (CV = 60 cm/s) at different pixel pitch values (representing the different FOV across the different species and tissue preparations) for the recommended (default) CV estimator engine (Polyfit). Typical operating range (jitter measured as the residual of a local quadratic fit to the activation maps of the 15 validation recordings) for conditioned signals in rat/rabbit data sets (0.09-0.24 ms), and wedge (0.45-2.0 ms) shown by the stippled and hatched regions. **Fig. 7f** shows the plot of DF recovered by CADENCE vs pacing stimulus (ground truth) from recordings (88 recordings from 83 hearts) from different species (and pacing rates). The median absolute error was 0.03% (0.002 Hz), with 95^th^ percentile at 0.044 Hz, and 100% within half an FFT bin (0.122 Hz) over the 84 recordings with 1:1 capture (from 80 hearts). The lower plot is an error plot (DF – pacing rate) against the ground truth (pacing rate). The human data are LV wedge only and 2 points took their truth from the annotated pacing rate (instead of pacing stimulus which was not available).

**Fig. 7.**
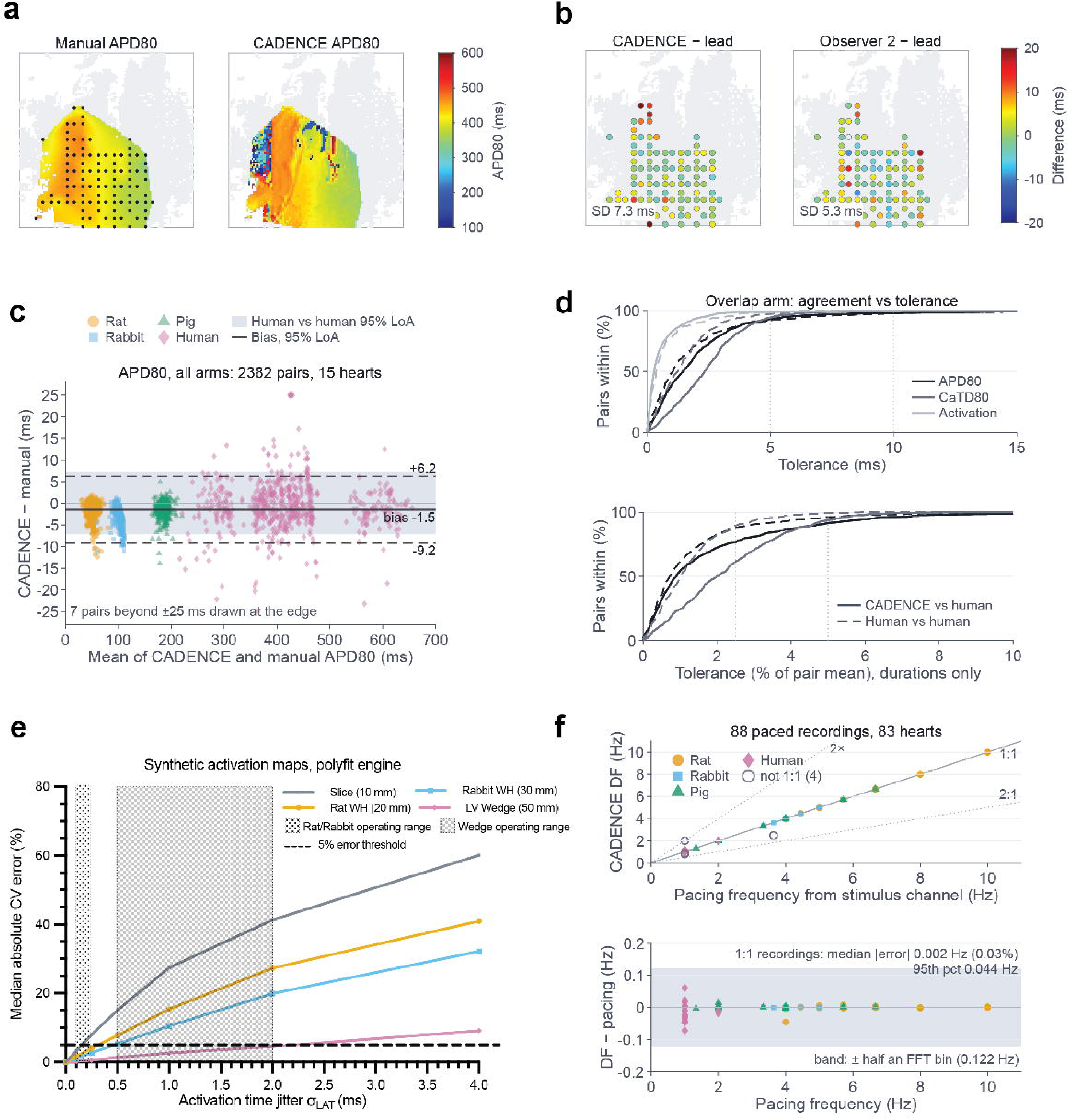
CADENCE validation: **a)** Mean APD80 map of the three observers (interpolated) and CADENCE APD80 map from an overlap human LV wedge recording (paced at 1000 ms). Black dots reflect location of pixels marked by the observers. **b)** Per pixel differences for the same APD80 pixel locations between CADENCE and lead observer (built the pixel pool), as well as between observer 2 and the lead observer. **c)** Plot of the Bland-Altman analysis with replicate measurements per subject (heart) without repeated measures correction. Pixels (APD80) are nested within recordings, observers, and hearts. Consequently, the pixel pairs (2382 pairs, 15 recordings, 3 observers) are not independent, and the pooled statistics are descriptive rather than inferential. **d)** Plot of the agreement vs tolerance between CADENCE vs human and human vs human for the marked metrics (activation time, APD80, and CaTD80). The lower plot uses a tolerance relative to the pair mean for APD80 and CaTD80 metrics. Both plots represent the overlap arm. **e)** Plot of median absolute CV error (%) vs activation time jitter computed from synthetic activation maps (CV = 60 cm/s) at different pixel pitch values for the recommended CV engine (Polyfit). Jitter was injected independently into each pixel’s activation time to generate a corrupted activation map. Typical operating range for conditioned signals in rat/rabbit data sets (0.09-0.24 ms), and wedge (0.5-2.0 ms) shown by the stippled and hatched regions. The bold dashed line represents the 5% error threshold in computing CV. The wedge is 100 × 100 pixels, and the others are 256 × 256 pixels. **f)** Plot of the DF recovered by CADENCE vs Pacing stimulus (ground truth) from recordings (88 recordings from 83 hearts) from different species (and pacing rates). The lower plot is an error plot (DF – pacing rate) against ground truth (pacing rate). The human data are LV wedge only and 2 points took their truth from annotated pacing rate (instead of pacing stimulus which was not available) and are shown as open pink diamonds. The open circles are 4 recordings that are not 1:1 capture. The gray band is ± half an FFT bin, 0.122 Hz.

## DISCUSSION

In this paper, we have presented CADENCE, an open-source MATLAB-based software platform with a modular, flexible architecture for end-to-end cardiac optical mapping data analysis. We demonstrated some of the CADENCE capabilities including signal conditioning, feature extraction and analysis, including action potential duration, voltage and calcium alternans, phase/rotor mapping, and conduction velocity across a range of optical imaging datasets spanning multiple species (rat, rabbit, pig, and human), multiple modalities (transmembrane potential, calcium transient), multiple tissue preparations (cardiac slices, wedge, and whole heart), and multiple camera configurations (single, dual, and panoramic).

CADENCE has been built on the research work done by various groups over the years^1,7,22,27,28^. We have incorporated key features of Rhythm, KairoSight, and ElectroMap^14^ in a single package (as shown in **Table 1**). Rhythm has robust signal conditioning capabilities including spatial binning, temporal filtering, drift correction, data normalization, and ensemble averaging. CADENCE builds on the same approach used in Rhythm to support signal conditioning features. Importantly, CADENCE has added support for new features that are not available in the other toolkits, including algorithms for SVD spatiotemporal denoising and motion correction. While motion correction has been implemented, we have not demonstrated the functionality in this study. CADENCE also provides support for panoramic (multi-camera) optical mapping data analysis like Rhythm, but not available in the other software toolkits.

CADENCE supports extraction of electrophysiology metrics from action potential (activation, repolarization, rise time etc.,) and calcium transients (decay time, duration, decay constant etc.,) like the other toolkits. KairoSight features image registration (for alignment of voltage and calcium signals), SNR based data curation, and in-depth alternans assessment, which CADENCE has also incorporated. ElectroMap has well-developed support for conduction velocity evaluation, which CADENCE has included in the released version. While Rhythm and ElectroMap both support arrhythmia analysis with display of phase maps, CADENCE supports tracking of annotated stable rotors and wavefronts.

We also demonstrated manual validation (CADENCE vs human) of metrics including APD80, activation time, and CaTD80, synthetic validation of CV, as well as the ability to accurately recover DF. The overlap arm entailed pooled, quantitative result across 1187 pairs (CADENCE vs human) for APD80 and activation time, and 597 pairs for CaTD80. For one observer pairing against the lead (using the second observer for comparison), the per-species APD80 SD (in ms) was human (7.3 vs 5.3), rat (0.94 vs 1.46), pig (1.08 vs 1.63), and rabbit (1.11 vs 0.83). The larger SD spread for human can be attributed to longer APDs compared during the overlap arm of the validation study. For the pooled, all observer pairings, the SD ratios (CADENCE vs human divided by human vs human) for APD80, Activation time, and CaTD80 were at 1.07, 0.95, and 0.89 respectively (as shown in **Supplementary Table S1**). Per-species APD80 SD ratios were for human 1.04, pig 0.82, rabbit 1.57, and rat 0.81. Activation times (LoA -2.6 to 2.9 ms) agreed far better than APD80 (LoA -9.2 ms to 6.2 ms) as the activation is a single steep upstroke crossing (50%), whereas APD80 depends on a repolarization level that is genuinely ambiguous on noisy signals, which is then associated with relatively higher CADENCE vs observer disagreements.

We also validated CV metrics using synthetic data (as observers can’t hand-mark CV) by generating activation maps (at different pixel pitch values) for isotropic planar wavefronts as well as for anisotropic conduction. For isotropic conduction, all three estimators (Gradient, Polyfit, and Circle) computed the expected CV (velocity range 20-100 cm/s). In case of anisotropic conduction, longitudinal and transverse CV was recovered with a field median absolute error of 0.17% or less on 256 x 256 grids and 0.88% or less on 100 x 100 wedge. Moreover, the anisotropy ratio was recovered within 1.2% for gradient, 2.6% for Polyfit and 3.9% for the Circle CV estimator. In case of jitter (activation timing precision), all three engines were able to recover CV close to the 5% CV error threshold for rat and rabbit conditioned signals at the measured activation jitter (0.1-0.25 ms). For human LV wedge conditioned signals, at the measured activation jitter (0.45-2.0 ms), the CV error was within 5% for Polyfit (4.5%) and Circle (4.6%) estimator but exceeded the threshold for Gradient (8.0%). However, for pig slice conditioned signals, at the measured activation jitter (0.8-2.1ms), the CV error was about 22-41% for the Polyfit and 37-55% for the Gradient (see **Supplementary Table S2-S5**, **Supplementary Fig. S2**). However, the Circle estimator broke down (aggressively discarding pixels at higher rates). While all three CV estimators were competitive, Polyfit was the recommended option for CADENCE as it was more robust across the full jitter range (0.1-2.0 ms), compared to the Circle estimator, which broke down depending on pixel size (e.g., at 0.5 ms for slice with 31.8% error, at 1 ms for rat whole heart with 32.1% error, and 2 ms for rabbit whole heart with 55.2% error). Gradient engine ran about 1.3-1.9x Polyfit’s error throughout.

Finally, we validated DF metrics using 88 recordings from 83 hearts across different species (and pacing rates). CADENCE was able to recover DF with high degree of fidelity (median absolute error was 0.03% (0.002 Hz) for recordings with 1:1 capture). The quantitative validation of CADENCE should offer a degree of confidence in using the software toolkit for optical mapping data processing and analysis.

CADENCE does come with a few limitations. First is the ability to reliably extract APD80 and/or CaTD80 at fast pacing cycles (85 ms or faster). This is not a software defect but rather a constraint associated with fewer pixels crossing the 80% repolarization threshold. Consequently, CADENCE recommends using APD50/CaTD50 at faster pacing cycle lengths. Second, computing CV anisotropy ratio (L/T) requires a non-planar wavefront. As such, CADENCE uses a plane-fit R^2^ > 0.8 gate because the edge-paced or planar sweep otherwise can produce spurious anisotropy ratios. Third, computing CV from tissue with small FOV (e.g., cardiac slice with 10 mm) is not reliable as CV error scales with jitter divided by pixel size (smaller pixel pitch results in larger CV errors). Fourth, CADENCE was designed to support end-to-end pipeline for cardiac optical data processing and analysis. However, the current release supports only two imaging modalities (transmembrane potential and calcium). Other modalities such as NADH and FAD are not yet supported^29^. Fifth, while CADENCE can display annotated rotors and wavefronts, additional metrics relative to wavefront dynamics (wavefront repeatability, collisions, fragmentation etc.,), and rotor dynamics (rotor displacement and trajectory, etc.,) are not currently available in this version of CADENCE. However, we are working on the next update to CADENCE that will include a separate arrhythmia analysis module, which will allow the user to extract and visualize these metrics. Another planned update for CADENCE includes AI-based feature detection and machine learning module for predictive metrics including for arrhythmia substrate assessment, and rotor localization for electrotherapy.

CADENCE was developed using MATLAB and is distributed as an open-source package with freely available code, it is also available as a compiled standalone executable that the end-user can install and run without needing MATLAB License. This will allow researchers without computational background to use CADENCE with minimal technical constraints. CADENCE software platform is also setup as a community contribution model via GitHub, to enable the research community to interact with, contribute to, and maintain the source code. We hope this will facilitate wider adoption and utilization of CADENCE software and potentially pave the way for standardization across research labs.

## DATA AVAILABILITY

The source code and documentation for CADENCE software is available for download from https://github.com/kedararas/Cadence (DOI: 10.5281/zenodo.23067078). Additional datasets are available from the corresponding author on reasonable request for the purposes of developing new methods of analysis, reproducing the results or replicating the procedure. Detailed description of the materials and methods are available in the supplementary materials.

## ETHICS STATEMENT

The animal study was reviewed and approved by Institutional Animal Care and Use Committee of University at Buffalo. Studies using de-identified human heart tissue were reviewed and approved by the Institutional Review Board at University at Buffalo.

## AUTHOR CONTRIBUTIONS

KA, AA, SP, EM, AL, NQ, and LB performed the experiments and approved the manuscript. KA, AA, SP, EM, AL analyzed the data. KA, AA, and SP prepared the figures. KA, AA, and SP drafted the manuscript. KA, AA, SP, EM, and AL conceived and designed the experiments. All authors contributed to the article and approved the submitted version.

## ACKNOWLEDGEMENTS

The authors thank Freddy Huff, Nathan Conrad, and Sam Verratti for their excellent technical assistance.

## SUPPLEMENTARY MATERIAL

The Supplementary Material for this article can be found online.

---

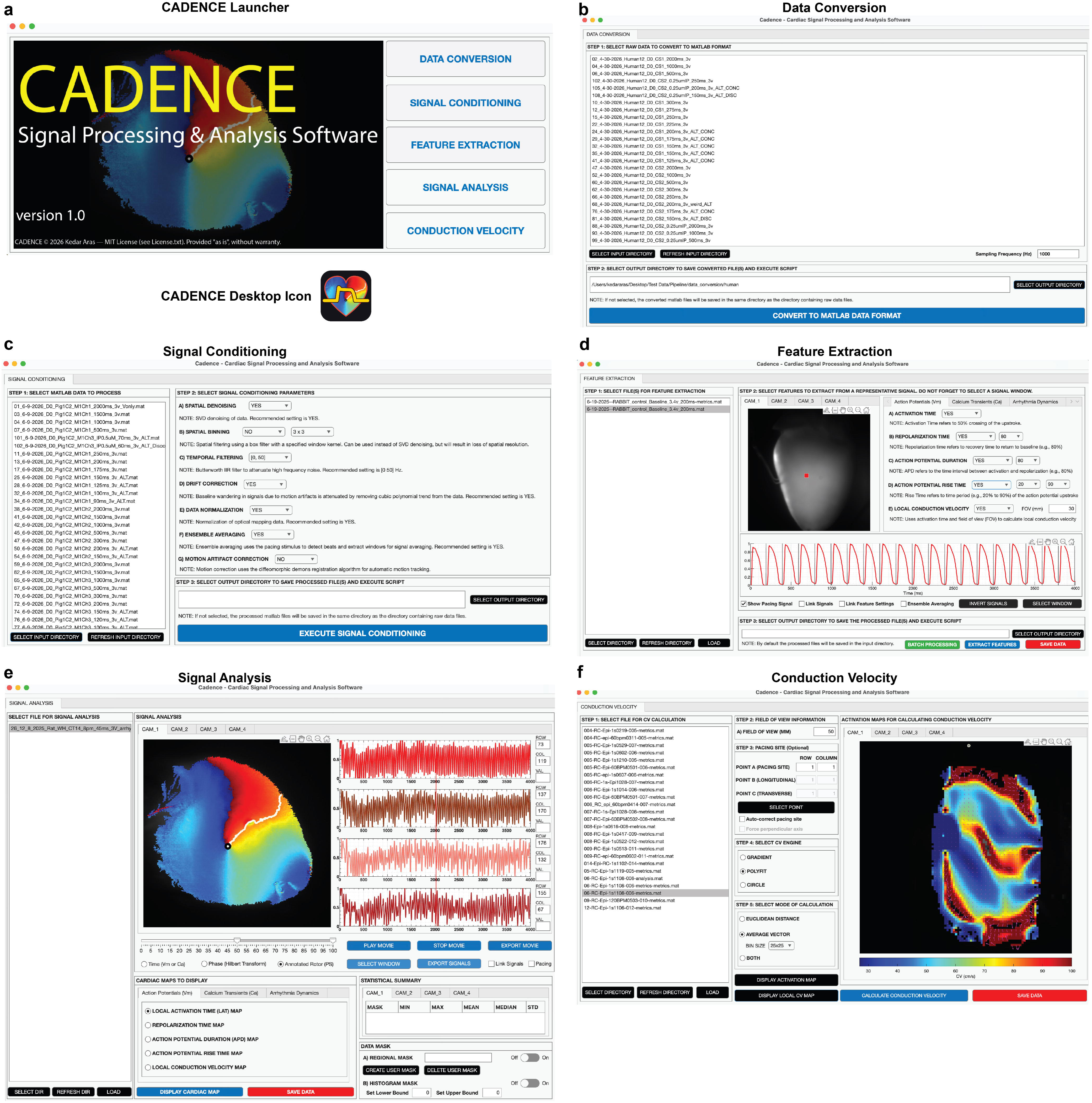

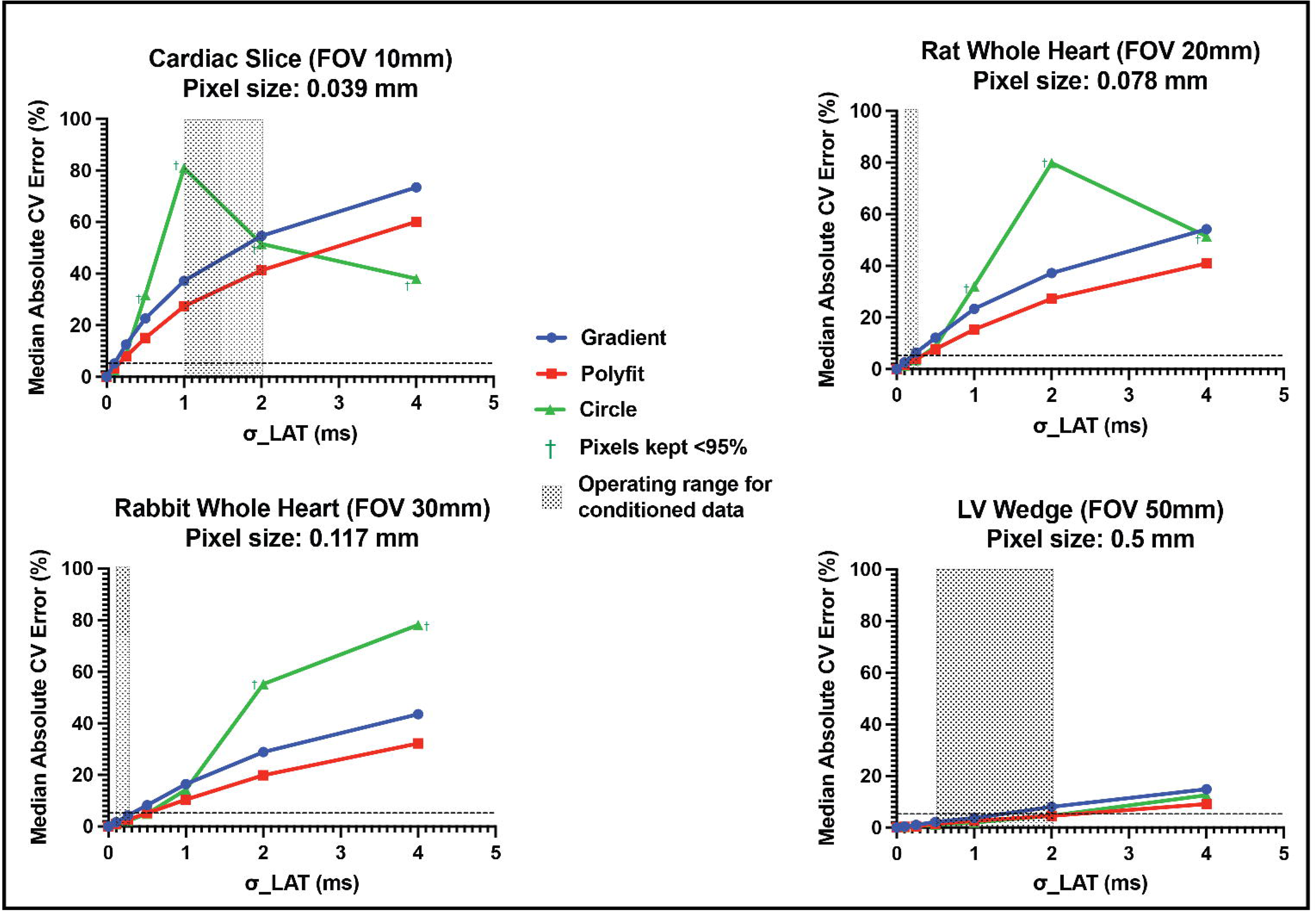

