## Supplementary Information for "CADENCE: Open-source software for signal conditioning, feature extraction, visualization and analysis of cardiac optical mapping data"

### DETAILED METHODS

#### System Requirements

The system requirements for running CADENCE software include MATLAB Runtime (2025b) (https://www.mathworks.com/products/compiler/matlab-runtime.html). CADENCE source code execution in MATLAB requires Signal Processing, Image Processing, Statistics and Machine Learning, and Curve Fitting. Currently supported for MacOS on Apple Silicon, and Windows. The minimum hardware requirements include 16GB RAM, 4-core 64 bit CPU, 50 GB SSD free storage, and 1920 x 1080 display. The source code and documentation for CADENCE software is available for download from https://github.com/kedararas/Cadence (DOI: 10.5281/zenodo.23067078).

#### Data Acquisition

All studies using animal models including pig, rabbit, and rat followed relevant guidelines and regulations that were approved by the Institutional Animal Care and Use Committee respectively at University at Buffalo (202200063, 202200066, 202300053). Use of de-identified human heart tissue (Connect Life, Buffalo, NY) was approved by Institutional Review Board (University at Buffalo). All procedures followed the National Institutes of Health Guide for the Care and Use of Laboratory Animals. For this study, we conducted optical mapping studies on 4 rat (Sprague Dawley, female, 8 weeks), 4 rabbit (New Zealand white, female, 3 months, 4 lbs.), 4 pig (Yorkshire, female, 70 lbs., 3 months), and 4 donor human hearts.

Donor human hearts were procured from ConnectLife (Buffalo, NY). All hearts were arrested using ice-cold cardioplegic solution in the operating room and transported to the laboratory for dissection and subsequent optical imaging studies. Pigs were pre-medicated with telazol and xylazine administered via intramuscular (IM) injection. Butorphanol was given as an added analgesic administered IM. Surgical plane of anesthesia was then induced and maintained with isoflurane. Through a midsternal thoracotomy, the heart was exposed and potassium chloride administered (intracardiac) to induce cardiac arrest. The heart was then explanted and flushed with ice-cold cardioplegic solution like donor human hearts. Rabbits were sedated using a ketamine and xylazine cocktail administered IM. Once sedated, a cocktail of heparin and sodium pentobarbital was administered IV using the ear vein to induce cardiac arrest. The heart was then excised using a sternal thoracotomy and the aorta cannulated for subsequent Langendorff perfused whole heart optical imaging studies. Rats were anesthetized using isoflurane. Once the reflex test failed to elicit a response, the heart was excised and the aorta cannulated like with rabbits.

To validate CADENCE software across a variety of optical imaging datasets, the experimental setups included single camera ($V_m$ or Ca), dual camera ($V_m$ and Ca) and panoramic (four cameras) configuration. We also employed three different tissue preparations including: cardiac slices (pig), left ventricular (LV) wedge (donor human heart, pig), and whole heart (rat and rabbit). Cardiac slices were sourced from the LV basal anterior or posterior regions of the heart. The tissue block from the LV was mounted on an agarose-coated specimen holder using surgical glue with the epicardial surface facing down. The specimen holder was placed in a cold bath (Tyrode's solution) of a high precision vibratome (7000smz-2, Campden Instruments). The tissue block was sliced using a ceramic blade, longitudinal to the myocardial fiber orientation and from endocardium down to the epicardium to produce the cardiac slices (350μm thickness). For wedge tissue preparations, transmural wedges from the posterolateral LV free wall supplied by the left marginal artery were isolated. Each wedge contained a section of coronary artery along its length, which was then cannulated for subsequent perfusion and optical mapping studies.

Cardiac tissue slices were mounted in a temperature controlled (37° C) bath containing oxygenated Tyrode solution[1]. Similarly, cannulated tissue preparations (Whole heart or LV wedge) were maintained in a temperature controlled (37° C) Langendorff perfusion system with oxygenated Tyrode solution at constant perfusion pressure of 60-80 mm Hg[2-4]. Tissue was immobilized with blebbistatin (5-15 μM, Cayman Chemical) to suppress motion artifacts in optical recordings, without adverse electrophysiological effects. A bipolar pacing electrode was used to electrically stimulate the tissue. For dual imaging (Vm and Ca) experiments, the tissue was loaded with Rhod-2 AM (calcium sensitive dye) and RH237 (voltage sensitive dye) and then illuminated using 520 nm excitation light source and the emitted (700 nm) voltage and calcium (585 nm) signals were recorded using two MiCAM03/05 (SciMedia, CA) CMOS cameras (256 x 256 pixels or 100 x 100 pixels, 1kHz sampling rate). In case of transmembrane potential ($V_m$) only optical imaging experiments, the tissue was loaded with Di-4-ANEPPS or Di-4-ANBDQBS voltage sensitive dyes. The optical action potentials were then recorded from up to four MiCAM03/05 CMOS cameras at high spatial and temporal resolutions. Study protocol entailed conducting standard S1-S1 pacing restitution protocol at decreasing pacing cycle length until loss of 1:1 capture. We then administered isoproterenol (0.5-1 μM) to shorten action potential duration and thus create a substrate for arrhythmia. Arrhythmia was then pace-induced using S1-S1 protocol or 1-second burst pacing at 60Hz.

### Software Architecture

**Fig. 1** depicts the CADENCE software architecture that follows a modular design philosophy with independently usable modules including for data conversion, signal conditioning, feature extraction, signal analysis, and conduction velocity. Graphical user interfaces (GUI) for each module were created using MATLAB App Designer, which employs the object-oriented coding framework (see **Supplementary Fig. S1**). Importantly, the Model-View-Controller (MVC) software design pattern compartmentalizes the user interface code from the back end algorithms and thus enables isolating software features into separate maintainable components and simplifies the process of adding new features to the software toolkit.

**Supplementary Figure S1:** CADENCE Software Modules **a)** CADENCE launcher and desktop icon **b)** Data Conversion module **c)** Signal Conditioning module **d)** Feature Extraction module **e)** Signal Analysis module **f)** Conduction Velocity module

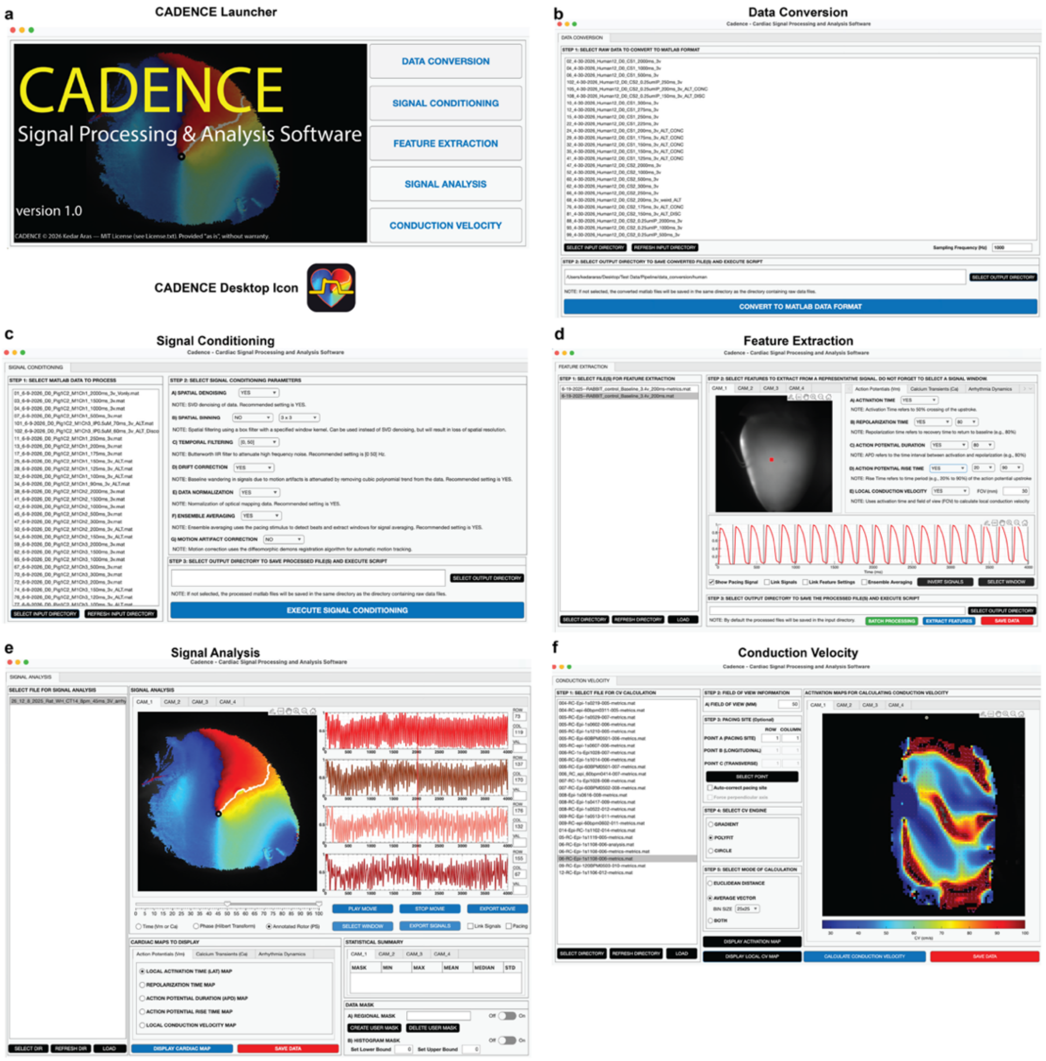

Data Import and Conversion Module: The first step in processing optical imaging data using CADENCE entails converting raw data files into MATLAB file format. The module allows the user to load TIFF or SciMedia format (*.gsh/*.gsd) raw data files for conversion to MATLAB (.mat) format. Typically, an optical mapping study entails acquiring multiple data recordings. Consequently, the module allows the user to select multiple data files for batch processing.

Signal Conditioning Module: The signal conditioning module allows the raw data files already converted into MATLAB file format to be processed to boost signal quality by filtering out noise. The signal conditioning pipeline comprises, in order: generating SNR map, drift correction, spatiotemporal denoising using singular value decomposition based low-rank decomposition [5] (falling back to 5x5 binning on any camera SVD cannot be applied to), spatial binning, temporal filtering, motion artifact correction[6,7], normalization, polarity check, ensemble averaging[8], and SNR mask (fixed threshold SNR $\geq$2). While motion correction algorithm is implemented, we did not demonstrate it here as our data acquisition experiments used a mechanical uncoupler.

SNR map is computed on raw, un-detrended data. Per-pixel SNR is computed as the peak-to-peak amplitude of temporally smoothed signal divided by the standard deviation of the residual high-frequency (raw minus a centered moving average, window approximately 1% of the recording). Since the noise is estimated from the residual rather than from a designated diastolic window, the measure does not require an identifiable diastolic interval and remains usable at fast pacing rates and during alternans or arrhythmia.

Drift correction is implemented using MATLAB's in-built detrend function that fits a third-order (cubic) polynomial to every pixel's time course by least squares and subtracts it.

Optical mapping data (action potential and calcium transients) are highly correlated across neighboring pixels and successive beats, unlike noise, which is spatially incoherent. Consequently, coherent signal occupies a low-dimensional subspace of the recording, whereas noise is distributed across many low-variance dimensions. Spatiotemporal denoising using truncated SVD (with fixed rank of 8)[5] takes advantage of this separation, and attenuates noise while preserving spatial resolution, unlike spatial binning, which trades resolution for signal quality.

As an optional alternative to SVD denoising, spatial binning (off by default) raises the signal-to-noise (SNR) ratio by averaging each pixel with its neighbors, at the cost of spatial resolution. An N x N uniform (box) average is applied to each frame implemented as convolution with an N x N kernel of ones and is computationally cheaper compared to SVD denoising.

High-frequency acquisition noise is attenuated with a fourth-order Butterworth infinite-impulse response (IIR) low-pass filter, implemented in second-order-section form and applied forward and backward along the time axis of every pixel.

CADENCE implements marker-free, two-stage tracking scheme for motion correction (off by default). First, every frame is locally contrast normalized – the local mean is subtracted and residual divided by a local standard deviation, which suppresses the slow, low-spatial frequency fluorescence changes of action potential, while preserving the fixed tissue texture (e.g., trabeculae), that moves with the myocardium. Second, a dense non-rigid displacement field is estimated from each frame to a single reference frame by multi-resolution diffeomorphic demons registration. The field is computed on the normalized video but applied to the original fluorescence frames. Motion correction is computationally expensive, especially over long recordings, and is intended for working heart optical mapping data, where uncoupling is not used.

For visualization and cross-pixel comparison, each pixel's time course is rescaled to [0, 1] using robust percentile bounds (1st and 99th percentiles) rather than min-max scaling, so that a single anomalous sample (e.g., filter edge artefact) does not define the range and compress the true signal.

Polarity check of signals (masked pixels with SNR $\geq$ 3) relies on the stimulus anchor and for every stimulus (which counts as a beat), the signal rise (max(beat) – base) and fall (base - min(beat)) is computed. After dropping the first and last beats, the median of each across beats (minimum 5 beats required) is used to compute confidence (-1 to +1), with negative suggesting inverted signals. If absolute confidence is < 0.2, or pacing stimulus is unavailable, the masked pixels are pooled into one trace weighted by SNR, histogram mode (correctly oriented signals spend most of their time near rest), and slope asymmetry (upstroke is faster than repolarization), which is given additional weight (counted twice). The inverted signals thus determined are flipped and per-recording confidence saved in the data file for provenance. Additionally, both checks now save a signed value, and the source check is also recorded.

When cardiac tissue is paced, repeated beats are near-identical and can be averaged to raise SNR without requiring temporal smoothing. Beat onsets are taken from pacing stimulus or detected from optical signal, when no stimulus trace is available. Each beat's baseline is subtracted, and outlier beats are rejected per pixel (RMS deviation above median + 3 $\times$ max(1.4826•MAD, 0.10•median), but only when there are 4 or more beats in a trace. The beats are averaged frame-by-frame and normalized to yield one representative beat per pixel. Finally, tissue masks are then formed by a pre-defined SNR floor (fixed threshold SNR $\geq$2).

Feature Extraction Module: Pre-processed optical mapping data can be loaded into the feature extraction module for post-processing and extracting relevant cardiac electrophysiology metrics. The post-processing features include data masking and image alignment (in case of dual voltage and calcium imaging). The user interface for data masking supports data curation based on region of interest (ROI) defined by a user created mask and/or mask based on signal-to-noise ratio (SNR) threshold. Adaptive SNR masking is a key feature of feature extraction and uses the rule SNR threshold = max (1.5, 0.5 x median SNR of pixels with SNR > 1.5).

CADENCE supports image alignment by registering the moving camera image (whichever is active, calcium or voltage) onto the fixed reference (e.g., voltage or calcium) and applying the resulting transform to the whole moving video stack. Because both cameras image the same tissue at the same pixel size, the default geometric model is a rigid transform (translation and rotation only). Registration is intensity-based using a gradient-descent optimizer (MATLAB 'imregtform', monomodal configuration). As with signal conditioning, the transform is estimated on the masked (relaxed Otsu threshold, hole-filling, and morphological closing), and contrast is enhanced with CLAHE.

Action potential metrics that can be extracted using CADENCE include local activation time (LAT), repolarization time, action potential duration (APD), and rise time. Similarly, calcium transient metrics that can be computed include voltage and calcium delay (for dual camera only, CAM1 = $V_m$ , CAM2 = Ca), decay time (CaDT), transient duration (CaTD), decay constant ($\tau$ ), and rise time. LAT is defined per pixel as the instant at which the optical upstroke crosses 50% of its baseline-to-peak amplitude, estimated with sub-frame resolution. For each pixel, the baseline and the peak of the beat are taken as the trace minimum and maximum, the threshold set at their midpoint, and the crossing time located by linear interpolation between the two frames that bracket it near the point of maximum dV/dt. If the steepest segment sits below 50%, the crossing is extrapolated. The map is also despeckled at 5 ms (pixels more than 5 ms from their 3 x 3 neighborhood median are replaced by it). Sub-frame interpolation removes the quantization of frame-limited timing and improves the accuracy of the downstream conduction velocity calculation. Repolarization time is the interval from start of the analysis window to the specified level of repolarization (default 80%). For a normalized post peak trace, the first time the signal falls to or below the level L (referenced to each pixel's pre-activation baseline and post-activation peak) is found and refined by linear interpolation between the bracketing samples. APD and CaTD at a given percentage are the intervals from activation to that repolarization/reuptake level (e.g., APD80, CaTD80). The same interpolated-crossing routine applied to calcium channel yields the calcium reuptake time (CaDT). The calcium decay constant ($\tau$ ) is obtained as a single exponential fit after baseline subtraction: a weighted log-linear seed refined by damped Gauss-Newton, with amplitude gate and a secondary estimate that relies on the crossing at the 1/e level of the post-peak transient. The voltage-calcium activation delay is the per-pixel difference between the calcium and voltage activation times. The above metrics are computed using the ensemble-averaged beat unless the user selects a representative beat for feature extraction. Using ensemble-averaged beat is ticked off by default.

CADENCE uses full beat series to extract alternans and arrhythmia analysis. Alternans is quantified beat-to-beat at every pixel.  CADENCE supports 15 different alternans metrics including: APD30/50/80/90 alternans, AP amplitude alternans, Upstroke dV/dt alternans, Triangulation (APD80 – APD30) alternans, Diastolic $V_m$ (load) alternans, AP release map, CaTD50/80 alternans, CaT amplitude alternans, Time-to-peak alternans, and Diastolic Ca (load) alternans map (diastolic difference over the larger beat's amplitude: D/L), and Ca release map[9]. Except for the Triangulation, AP and Ca load maps, and AP and Ca release maps, the remaining 10 alternans metrics have a corresponding dimensionless ratio map (|mean(odd-even)|/mean), which is the same measurement normalized. Alternans phase (APD80 phase and phase angle, AP amplitude phase, and CaT amplitude phase) is also computed with the binary phase being sign(mean(odd-even)) representing APD80 for voltage, and amplitude for calcium. The FFT at alternans bin gives the separate continuous phase angle. Alternans is classified as spatially concordant or discordant based on the sign of the APD alternans at the substrate's automatic level on significant pixels, meaning alternans above 15% of mean APD. If the resulting map has majority fraction under 0.8, the alternans is deemed discordant. Per pixel significance including spectral alternans index, paired t, and p-value is also supported for both $V_m$ and Ca. Finally arrhythmia substrate is characterized using Ca-AP coupling, concordance, nodal lines, APD gradient, diastolic Ca, and composite risk score. In this study, we demonstrate some of the alternans metrics, but they are not validated (compared against manual marking).  During APD and CaTD metrics extraction, if the resulting values are non-finite (NaN or ≤0), they are excluded from analysis and are associated with holes (gray regions) shown in the cardiac maps. As seen in **Fig. 4**, at 85 ms pacing cycle length, CaTD80 is reached in only 47.8% of tissue beats.

For arrhythmia analysis, each pixel's power spectral density is estimated by Welch's method. The dominant frequency (DF) is the peak of the power spectral density (PSD) and uses the rate adaptive search band (set once per recording from the peak of the pooled spectrum over 0.5 to 60 Hz) and parabolic sub-bin peak refinement with the following Welch parameters: 2-second Hann windows, 50% overlap, NFFT = max(256, $2^{\text{nextpow2(4.fs)}}$), which is 4096 at 1kHz for 0.244 Hz bins; 4 harmonics (K); search band [max(0.5, 0.5• $f_0$), min(0.9 •Nyquist, 2• $f_0$)]; peak half-width max(0.5, 0.05 •DF); denominator [0.5, (K+0.5) •DF]. Regularity index (RI) is the fraction of in-band power concentrated in a narrow window about DF[10]; and the organization index (OI) additionally sums the power of DF and its harmonics[11], so that an organized action-potential train yields OI >> RI (holds only when there is harmonic power), while fibrillatory activity gives low, comparable values. For rotor analysis, each pixel's signal is converted to an instantaneous phase via the Hilbert transform, yielding a wrapped phase map on $(-\pi, \pi)$ per frame. Phase singularities (PS) are located by computing the topological charge at every pixel[3]. The resulting PS are tracked spatially (linking distance = 1mm of tissue converted to pixels) over time to identify stable rotors (rotors are considered stable if they last at least 1.5 times the rotation period, which uses $75^{th}$ percentile of the DF map). PS are matched to a velocity-extrapolated predicted position with the same charge sign, and not to the previous location. A PS may "vanish briefly" (only if the next frame contains some PS) and still link forward. The look ahead is min(half of a rotation, max(17 ms, quarter rotation)) and the distance threshold widens with $\sqrt{(\text{frames ahead})}$, capped at 3x.

<u>Signal Analysis Module:</u> Features extracted from the optical mapping data can be analyzed using the signal analysis module. The user interface supports spatiotemporal analysis of optical mapping data via display of two-dimensional (2D) cardiac maps, 3D movies including voltage or calcium transient, phase, and annotated rotor (which tracks wavefronts and stable rotors), and pixel-level action potential and/or calcium transient signals.

<u>Conduction Velocity Module:</u> Conduction velocity can be computed using the CV module, which accepts the local activation map (LAT) as an input. As described in the previous section, LAT is the timepoint corresponding to 50% of the base-to-peak upstroke amplitude and is computed by temporal linear interpolation between two bracketing frames. This was a conscious choice instead of using max dV/dt, as at small field of view (e.g., 10 mm) and high spatial resolution (e.g., 256 x 256 grid, pixel pitch 0.039 mm), the max dV/dt approach can create neighboring pixels with same activation time (resulting in divergent CV values) or edge pixels with sudden change in activation time step (e.g., by 1ms) and the resulting CV is too slow. The sub-frame linear interpolation is computed using the equation

$LAT = k + (t - v(k)) / ( v(k+1) - v(k))$ where

$v$ = pixel fluorescence trace

$t$ = crossing threshold, per pixel ($t = base + 0.5*amplitude$)

$base = \min(v)$ over the window

$peak = \max(v)$ over the window

$amp = peak - base$

k = last frame at or below threshold

v(k+1) = next frame, above threshold

The resulting LAT is computed in frames and converted to ms (x1000/acquisition frequency). Pixels where the threshold isn't bracketed take the integer max dV/dt frame as long as dt > 0. Moreover, pixels that do not meet the amplitude (>10% of max) or slope (>20% of the median) threshold are rejected (NaN assignment). LAT is not computed for pixels that lie outside the data mask (user selected and/or SNR threshold). The field of view is user-specified, which serves as the input for pixel pitch (mm/pixel). For CADENCE validation, we tested different pixel pitches across multiple tissue preparations including cardiac slice (0.039 mm, 256 x 256 grid), rat whole heart (0.078 mm, 256 x 256 grid), rabbit whole heart (0.117 mm, 256 x 256 grid), and human LV wedge (0.5 mm, 100 x 100 grid).

CADENCE offers two methods (Average vector and Euclidean) and two estimators (Gradient, Polyfit) to compute CV, which inverts the activation gradient using the formula

CV magnitude = $1/ ||\nabla T||$; CV direction = $\nabla T/ ||\nabla T||^2$;

Gradient engine employs the finite differences of the Gaussian smoothed (default $\sigma = 2$ pixels) activation map, whereas Polyfit engine uses the local least-squares surface fit (7 x 7 window, second order quadratic fit) [12]. In addition, CADENCE also supports the Circle engine, which samples 36 chords[13], and uses a default radius of 12 pixels (based on the bin size selector, which the user can change) then takes the direction with the largest $|\Delta LAT|$ and back-projects speed over a $\pm 60°$ wedge.

For the average vector CV calculation method, CADENCE supports both manual and automatic modes. In the manual mode, the user provides locations of A (origin), B (longitudinal), and C (transverse) sites, which is then used to determine local kernels centered on B and C to compute the CV. If number of qualifying pixels is less than 25 pixels, CV calculation uses the whole-field pooling within $\pm 30°$. In the automatic mode, a region of interest (ROI) centered on the origin site (user selected or auto-detected) of fixed radius (e.g., 0.25 * min(rows, cols)) used to compute longitudinal and transverse CV. Pixels are also accepted or rejected based on velocity cap (speed min > 1 cm/s, speed max = 250 cm/s, with pixels over the cap treated as NaN and below the minimum threshold excluded from pooling), and gaussian smoothing (NaN aware normalized convolution) kernel support (the output pixel at the kernel center is rejected, if the valid kernel fraction falls below 50%). Edge pixels (8 pixel erosion of the tissue boundary) are also excluded (applies to all directional pooling) because the inverse-gradient method inflates CV at boundaries. The total number of pixels used for computing CV is reported alongside CV summary for reference.

CADENCE employs user-selected pacing site (origin), if available. Alternately, it can auto-detect the earliest activation site (centroid of the earliest activated region) within 1ms of a robust 0.5th percentile floor, after despeckling and rejection of zero/negative times. The manual point (user selected) is overridden, if auto-selection option is enabled (off by default), and the selected pixel location differs from the auto-detected pixel location by more than 15 pixels or greater than 5 ms of local activation time. A region of interest centered on the origin site with a radius (0.25 x min (row, cols), e.g., for a 256 x 256 grid, ROI would be a disc of radius 64 pixel and encompass half of the field width) is selected and the intersection of the valid pixels and pixels within the ROI is used to compute the longitudinal and transverse CV. The choice of employing ROI instead of the whole field, was to minimize the dilution of the anisotropy associated with rotation of fiber orientation. However, if fewer than 50 pixels survive the ROI, the fit reverts to whole field instead and is reported along with the CV summary. The local CV magnitudes are binned in 15° bins ( ≥8 pixels per bin) and each bin contributes its median CV, and the least-squares problem has at most 12 data points, which is solved linearly in $V^2$ space. The fitted form is

$$V^2 = A + C \cdot \cos 2\varphi + D \cdot \sin 2\varphi$$

Then $L = \sqrt{(A + B)}$ and $T = \sqrt{(A - B)}$, fiber axis ($\theta$) = ½ ·atan2(D, C), $B = \sqrt{(C^2 + D^2)}$

For valid L and T values, the total number of pixels used for computing CV should be ≥ 50 pixels and ≥ 4 populated bins. Degenerate cases (e.g., where $T^2 < 0$) results in reporting max/min directional medians as L and T, with the axis taken from the maximal bin. The ellipse-fit $R^2$ describes the binned fit and explains the variance across the 12 directional medians. In addition, the ellipse-fit is weighted by pixel count. The reported values for L, T and $\theta$ are considered plausible if the ellipse-fit $R^2$ exceeds the threshold of 0.3 (can explain at least 30% of the variance). However, to ascertain the validity of L, T and $\theta$, we also assess planarity of the activation map. This is done by computing least-squares fit of a plane to the activation map itself. If the planar-fit $R^2 > 0.8$ threshold, the activation map is considered unidirectional (LAT increases linearly in one fixed planar direction at a constant rate) and the values of L/T are not reported. In this case, the speed along the fitted planes gradient is reported as the dominant CV (sweep CV) along with the wavefront direction (sweep direction), which is distinct from fiber axis ($\theta$). Beyond the ellipse-fit and the planar-fit, two additional guards are used to validate L, T, and $\theta$. First is the resolvability guard wherein, if more than 20% of valid pixels exceed 100 cm/s and the activation spread is less than 15 ms, then the ellipse-fit is deemed noise dominated and the L/T values are invalidated. Second is the cross-front artifact guard, if the fast axis (longitudinal CV direction) is within 20° of perpendicular to a strongly dominant propagation direction (circular concentration R > 0.5), then activation map is suspected to be planar, even if it passes the planar-fit threshold and the L/T values are deemed invalid. These four criteria (ellipse-fit, planar-fit, resolvability guard, and cross-front artifact guard) ensure strong quality check for unreliable L/T values.

Euclidean method of CV calculation relies on user selection of origin site (point A), longitudinal site (point B), and transverse site (point C). If not selected, the points A, B, and C are positioned at the auto-detected origin site, along the fiber axis, and at axis + 90° respectively. CV is then computed as a pure two-point arithmetic given as distance divided by activation time difference between the origin and the longitudinal or transverse point (as long as dt > 0). While the average method uses four quality check criteria for CV calculations as described above, the Euclidean method employs only the planar-fit $R^2$ guard to notify the user on the validity of the reported values.  Specifically, if the plane-fit $R^2$ check fails, the L/T values are reported with a warning, and the larger of L and T is displayed as the dominant CV. Moreover, the choice of estimator (gradient, polyfit, or circle) affects the Euclidean CV calculation only if the points B and C are auto positioned by CADENCE, which requires computing local CV field based on choice of CV estimator engine to determine the fiber axis.

**CADENCE Validation**

We employed blinded manual marking by three observers (15 hearts including 4 human, 4 rat, 4 pig, and 3 rabbit) for CADENCE vs human and human vs human validation. Briefly, the validation protocol comprised the overlap (4 recordings marked by all 3 observers) and breadth (each observer marking additional 4 recordings) arms. One rabbit recording was marked twice (breadth arm), so 16 different assignments cover 15 distinct recordings. For each recording, the observer marked diastolic baseline, activation time (50% crossing), peak, repolarization (80% crossing) for 100 curated pixels (signals deemed to be of good enough quality by the observer to successfully mark the four locations). The extracted metrics (activation, APD80, and CaTD80) were compared against CADENCE. In addition, 88 recordings (from 83 hearts including human 22, pig 15, rabbit 9, and rat 37) from datasets already available in our lab, were used to recover DF and compared against pacing stimulus (84 recordings had 1:1 capture). Pig wedges only contributed to the DF cohort. Moreover, human cardiac slices were excluded from validation dataset due to inadequate signal quality. The validation harness is shipped with the source code and was run in MATLAB 2025b.

**Statistical Analysis**

CADENCE validation is an agreement study as such no hypothesis study was performed. Comparisons are per-pixel pairs (CADENCE vs observer, observer vs observer) and includes 2382 pairs for APD80 and activation, and 1194 pairs for CaTD80. We used the Bland-Altman analysis with replicate measurements per subject without a repeated measures correction. Pixels are nested within recordings, observers, and hearts and consequently the pixel pairs are not independent, and the pooled statistics are descriptive rather than inferential. For agreement statistics, metrics include median and the 95$^{th}$ percentile absolute difference. Bias is defined as mean difference (CADENCE – observer), so a negative bias means CADENCE reads shorter. Standard deviation is calculated for the differences, and 95% limits of agreement is characterized as bias $\pm$1.96 SD. Pearson r is a descriptor, and not an agreement measure. Proportion of pairs within 5 ms and within 10 ms are reflective of 5 and 10 frames at 1 kHz. The SD is comparable as both comparisons share the same reference observer, so variance var(CADENCE – reference) = $\sigma^2$ CADENCE + $\sigma^2$ reference and var(observer – reference) = $\sigma^2$ observer + $\sigma^2$ reference (**Fig. 7b**). The SD ratio is computed for the overlap arm only and is defined as SD (CADENCE – each of the three observers)/ SD (all three observer pairs) with $\leq$1 suggesting CADENCE sits inside the interobserver spread. For comparison in the overlap arm, the reference observer selected 100 pixels per recording (ensemble averaged signals where marking is possible) from a candidate pool stratified by SNR tercile with a fixed seed, above the same adaptive SNR (ranged from 1.5 to 5.47 across pools) floor CADENCE feature extraction uses and oversampled by a factor of 2. Observers were blind to CADENCE's output and to each other's marks. For the breadth arm, CADENCE used the same pooled 100 pixels selected for marking by the observer for a given recording. CADENCE censoring rate (pixels a reviewer marked where CADENCE produced no value and were dropped from the analysis) was negligible (20/3596 = 0.56%). By comparison, the pooled (every pool builder) skip rate during pool construction was 39.8% (3990 candidates and skipped 1590). Thus, CADENCE validation claims agreement on pixels a trained observer judged markable, not on all tissue.

For DF analysis, the ground truth is the pacing stimulus channel. Error metrics include median absolute error (Hz), 95$^{th}$ percentile, maximum, bias, percentage within half an FFT bin (0.122 Hz), and as percentage of the pacing rate (e.g., within 1% and 2% of pacing rate). The accuracy of DF recovery was limited to 84 of 88 recordings, which had 1:1 capture (within the tolerance window w = max(5% of pacing rate, 0.55 x FFT bin width)). The other 4 recordings were dropped on account of 2 floor pinned (occurs when DF search band is auto-detected and more than half the fraction of tissue pixels has their peak sitting at the bands lower edge, suggesting that DF is not the real peak but edge of the search window), 1 2 x harmonic, and 1 other. The unit of analysis is the recording, and some hearts contribute more than one recording, which is why 88 recordings come from 83 hearts.

### Validation of APD80, Activation, and CaTD80

**Supplementary Table S1:** SD ratios per species for the validated metrics

| Species | Metric | CADENCE vs human | | | | Human vs human | | | | SD ratio (CADENCE vs human ÷ human vs human) |
|---|---|---|---|---|---|---|---|---|---|---|
| | | Pairs (n) | Bias (ms) | SD (ms) | Within 5 ms | Pairs (n) | Bias (ms) | SD (ms) | Within 5 ms | |
| **All (pooled)** | **Activation time** | **1187** | **0.21** | **1.19** | **98.8%** | **1198** | **0.23** | **1.25** | **99.1%** | **0.95** |
| **All (pooled)** | **APD80** | **1187** | **-0.67** | **3.95** | **92.9%** | **1198** | **0.05** | **3.69** | **91.8%** | **1.07** |
| **All (pooled)** | **CaTD80** | **597** | **-2.40** | **1.68** | **94.0%** | **600** | **0.88** | **1.88** | **97.2%** | **0.89** |
| Human | Activation time | 299 | 0.47 | 1.93 | 98.3% | 298 | 0.74 | 2.29 | 96.3% | 0.84 |
| Human | APD80 | 299 | 1.33 | 7.14 | 76.6% | 298 | -0.49 | 6.85 | 70.5% | 1.04 |
| Pig | Activation time | 300 | 0.29 | 1.26 | 97.0% | 300 | -0.03 | 0.64 | 100.0% | 1.98 |
| Pig | APD80 | 300 | -0.85 | 1.37 | 98.7% | 300 | 1.07 | 1.66 | 98.0% | 0.82 |
| Pig | CaTD80 | 300 | -2.45 | 2.06 | 89.7% | 300 | 1.33 | 2.25 | 94.3% | 0.92 |
| Rabbit | Activation time | 288 | 0.08 | 0.29 | 100.0% | 300 | 0.12 | 0.38 | 100.0% | 0.76 |
| Rabbit | APD80 | 288 | -1.27 | 1.15 | 99.7% | 300 | -0.34 | 0.73 | 100.0% | 1.57 |
| Rat | Activation time | 300 | 0.00 | 0.26 | 100.0% | 300 | 0.08 | 0.36 | 100.0% | 0.72 |
| Rat | APD80 | 300 | -1.91 | 1.41 | 97.0% | 300 | -0.02 | 1.73 | 98.7% | 0.81 |
| Rat | CaTD80 | 297 | -2.35 | 1.18 | 98.3% | 300 | 0.42 | 1.29 | 100.0% | 0.92 |

SD ratios are computed for the overlap arm only (the same curated pixels marked by all three observers ( 4 recordings: human LV wedge, rat, pig, rabbit). CADENCE vs human is defined as CADENCE minus each observer at each marked pixel. Human vs human refers to every observer pair at the same pixels.  Bias refers to mean difference, and SD is the standard deviation of the differences. Within 5 ms refers to share of pairs with absolute difference $\leq$5 ms. SD ratio < 1 suggests that CADENCE differs from the observers less than they differ from each other. Alternately, SD ratio > 1 indicates that CADENCE differs from observers more than they differ from each other.  Human and rabbit recordings do not have calcium data.

### Validation of Conduction Velocity Module using synthetic data

#### Test A: Isotropic Conduction

First, we built activation maps (256 x 256 Grid, pixel size: 0.039/0.078/0.117 mm; 100 x 100 Grid, pixel size: 0.5 mm) for isotropic planar wavefronts computed using the formula $T = (x \cos\theta + y \sin\theta)/v$, so that $||\text{gradient } T|| = 1/v$ and true CV = v at every pixel for any propagation angle, all three engines computed the expected conduction velocities (velocities: 20-100 cm/s, angles: 0-90°) without any intrinsic bias confirming the implementation verification as shown in Supplementary Table S2.

**Supplementary Table S2**: Isotropic (planar wavefront, per-pixel, closed-form truth)

| Engine | 20 cm/s (0 - 90°) | 40 cm/s (0 - 90°) | 60 cm/s (0 - 90°) | 80 cm/s (0 - 90°) | 100 cm/s (0 - 90°) | Median Error | P95 | Direction Error | Plane Fit $R^2$ |
|---|---|---|---|---|---|---|---|---|---|
| Gradient | 20 | 40 | 60 | 80 | 100 | 0.00% | 0.00% | 0.00° | n/a |
| Polyfit | 20 | 40 | 60 | 80 | 100 | 0.00% | 0.00% | 0.00° | 1.00 |
| Circle | 20 | 40 | 60 | 80 | 100 | 0.00% | 0.00% | 0.00° | n/a |

**Test B: Anisotropic Conduction**

Next, activation maps were generated for anisotropic conduction from a point source using the formula $T = \text{sqrt}((\xi/v_L)^2 + (\eta/v_T)^2)$, where $\xi$ = distance from the source along the fiber axis (mm) and $\eta$ = distance from the source across the fiber axis (mm). We then computed longitudinal and transverse conduction velocities along and across the fiber orientation respectively and local CV everywhere in between. All three engines were able to recover the longitudinal CV, transverse CV with a field median absolute error of 0.17% or less across engines and pairs, suggesting cross-engine agreement and internal consistency on identical maps as shown in Supplementary Table S3 for the 256 x 256 (cardiac slice, rat and rabbit) grid preparations.

**Supplementary Table S3:** Anisotropic (Elliptical wave, per-pixel, closed form truth) for 256 x 256 grid (Cardiac slice, Rat and Rabbit whole heart preparation)

| Cardiac Slice /Rat WH/Rabbit WH (256 x 256 grid) | | | | | | | | | |
|---|---|---|---|---|---|---|---|---|---|
| Engine | L60 | T25 | AR 2.40 | L80 | T32 | AR 2.50 | L40 | T20 | AR 2.00 | Field Median \|Error\| for 60/25 cm/s pair |
| Gradient | 60.1 | 25 | 2.41 | 80.2 | 32 | 2.51 | 40.1 | 20 | 2.00 | 0.02% |
| Polyfit | 60.3 | 25 | 2.41 | 80.4 | 32 | 2.51 | 40.1 | 20 | 2.01 | 0.03% |
| Circle | 60.4 | 25 | 2.41 | 80.6 | 32 | 2.52 | 40.2 | 20 | 2.01 | 0.16% |

However, for the 100 x 100 grid preparation (LV Wedge), the engines slightly overestimated the longitudinal CV, and the field median absolute error was 0.12-0.88%, suggesting the small positive bias seen at the 256 x 256 grid was amplified at lower resolution (Supplementary Table S4). This can be attributed to the engines smoothing over a fixed pixel window, resulting in averaging over a larger tissue area at coarser resolutions.

**Supplementary Table S4:** Anisotropic (Elliptical wave, per-pixel, closed form truth) for LV Wedge (100 x 100 grid)

| LV Wedge (100 x 100 grid) | | | | | | | | | | |
|---|---|---|---|---|---|---|---|---|---|---|
| Engine | L60 | T25 | AR 2.40 | L80 | T32 | AR 2.50 | L40 | T20 | AR 2.00 | Field Median \|Error\| for 60/25 cm/s pair |
| Gradient | 60.7 | 25 | 2.43 | 81.0 | 32 | 2.53 | 40.3 | 20 | 2.02 | 0.12% |
| Polyfit | 61.5 | 25 | 2.46 | 82.1 | 32 | 2.56 | 40.7 | 20 | 2.03 | 0.21% |

| | | | | | | | | | | |
|---|---|---|---|---|---|---|---|---|---|---|
| Circle | 62.2 | 25 | 2.49 | 83.2 | 32 | 2.60 | 41.0 | 20 | 2.05 | 0.84% |

**Test C: Jitter (Activation timing precision)**

For this test, we started with a synthetic activation map with true CV of 60 cm/s at every pixel location. Next, we injected noise (zero-mean gaussian noise of known standard deviation) independently into each pixel's activation time to generate a corrupted activation map. The three engines then used the corrupted map to recover the CV at each pixel for comparison against known truth. Supplementary Figure S2 shows the results plotted as median absolute CV error (%) against activation timing precision ($\sigma$_LAT) for the three engines across different pixel sizes (species and preparations). CV accuracy is affected by jitter and pixel size (error ≈ 2 x $\sigma$_LAT x v/ds, Polyfit at low jitter). When the activation map was not corrupted, the engines reported accurate CV values. CV sensitivity is adversely affected by pixel size, resulting in relatively increased median absolute error when working with smaller preparations (e.g., cardiac slices, FOV=10mm) in comparison with larger tissue preparations (e.g., LV wedges, FOV=50mm) on account of smaller per-pixel LAT step (0.065ms on a slice vs 0.833ms on a wedge). Importantly, the Polyfit and the Circle engines were able to recover CV within a field median absolute error margin of 5% across the operating ranges for rat, rabbit, and LV wedge as seen in Supplementary Table S5. The Gradient estimator engine exceeded that threshold (6.4% on rat, 8.0% on LV wedge).

**Supplementary Table S5:** Cross-engine performance in the operating range of conditioned data ($\sigma$_LAT = 0.1 ms – 2.0 ms)

| | Median Absolute Error (%) | | | |
|---|---|---|---|---|
| Preparation | Measured Jitter $\sigma$_LAT | Gradient Engine | Polyfit Engine | Circle Engine |
| Slice (10 mm) | 1-2 ms | 37-55% | 27-41% | 81% then 52%, discarding pixels |
| Rat WH (20 mm) | 0.1-0.25 ms | 2.6-6.4% | 1.6-4.0% | 1.3-3.5% |
| Rabbit WH (30 mm) | 0.1-0.25 ms | 1.7-4.3% | 1.1-2.7% | 0.9-2.2% |
| LV Wedge (50 mm) | 0.5-2.0 ms | 2.2-8.0% | 1.4-4.5% | 1.0-4.6% |

The Circle estimator broke down (aggressively discarding pixels at higher rates). While all three CV estimators were competitive, Polyfit was the recommended option for CADENCE as it was more robust across the full jitter range (0.1-2.0 ms), compared to the Circle estimator, which broke down depending on pixel size (e.g., at 0.5 ms for slice with 31.8% error, at 1 ms for rat whole heart with 32.1% error, and 2 ms for rabbit whole heart with 55.2% error). Gradient engine ran about 1.3-1.9x Polyfit's error throughout.

**Supplementary Figure S2:** CV Error vs Activation Time Precision (Cross Engine)

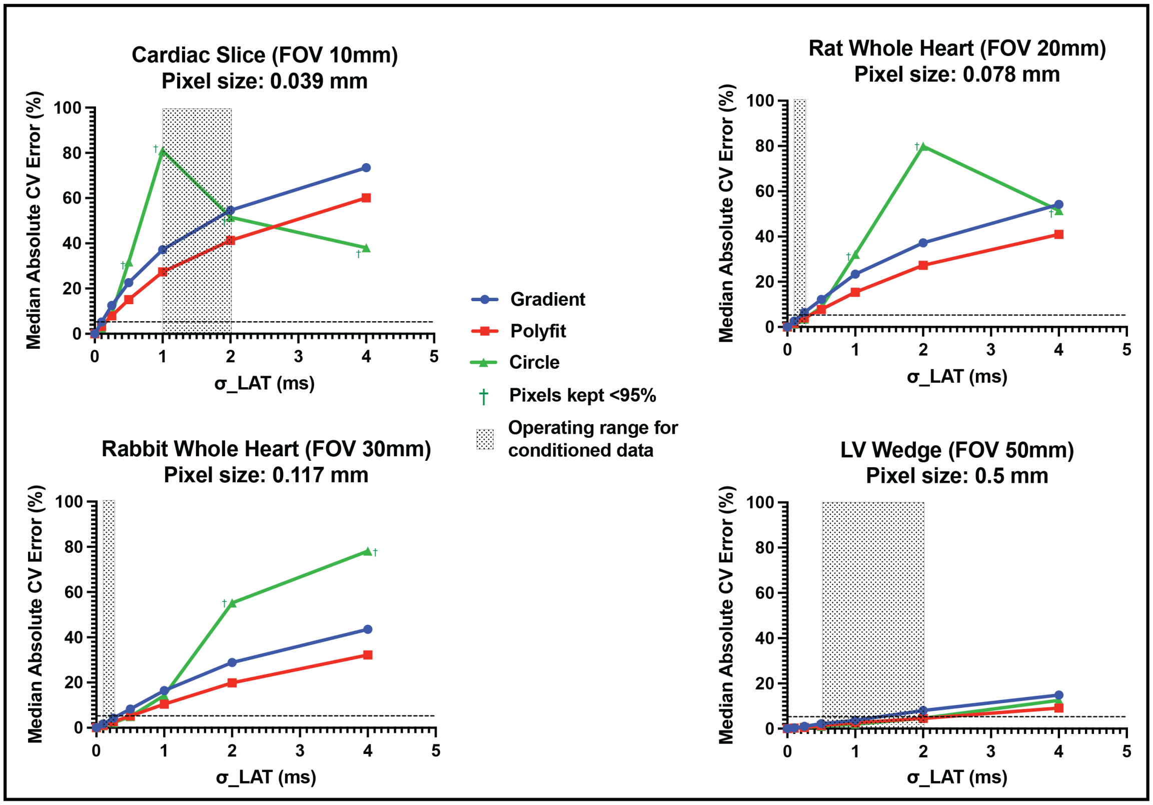


**Supplementary Video S1:** Example of monomorphic ventricular arrhythmia using a rat optical mapping data recording induced using S1-S1 pacing protocol with DF at 25 Hz. Rotors (black outline with white center) and wavefronts (white) are tracked. Every 5$^{th}$ frame of a 1kHz recording (230 ms total) is played at 10 fps with the clip duration at 4.7 s. The video shows the Hilbert phase with cyclic colormap. The non-valid (unstable) PS have gray-outline markers (with white center).
